# Two opposite failure modes in alignment-based conservation scores, one inflating and one concealing constraint, are computable before the score is used

**DOI:** 10.64898/2026.10.20.760363

**Authors:** Taner Karagöl, Alper Karagöl

## Abstract

A conservation score is read as a property of an amino acid position, when it measures that position together with a sequence set and a depth parameter, and the same position receives opposite calls under defensible choices of the latter two. We characterise two failure modes that push in opposite directions and show that both are computable from the alignment before the score is used. Where conservation is 1 − H/log 21 over an alphabet whose twenty-first symbol is the gap, a fully gapped column scores the maximum by construction. Across 55,816 columns the reported value rises with gappiness at ρ = +0.553, collapsing to +0.066 when conservation is recomputed over ungapped residues, and an independent predictor arbitrates: the gap-free recomputation tracks AlphaMissense at ρ = 0.551 against 0.416 for the reported value. The opposite mode is clade partitioning, in which between-subfamily divergence is read as tolerance. Varying only the comparator set moves a position by a per-family median of 2.19 grades on a nine-grade scale over 5,092 positions in eight families and flips 2.7 to 8.3 per cent, median 6.6, from variable to conserved. A label-permutation null cannot separate this from shared ancestry, calling 68.3 to 100 per cent of variable columns diagnostic; a tree-aware null with per-column parsimony rate matching returns 4.1 to 69.1 per cent, and the same four syndecan genes at two sampling depths give 4.1 and 29.8 per cent, so no proteome-wide rate is well posed without the decomposition behind it. The resulting per-position panel reports gap fraction, effective comparator count and the tree-corrected subfamily-explained fraction with the direction of the expected error, and generates a testable prediction, that variants to a sister subfamily’s fixed state are benign-enriched, which holds at an odds ratio of 2.46 (1.40 to 4.33) carried by 87 such variants among 1,565 classified. Applying the gap correction to a published conserved-score pipeline raises the rigid score’s discrimination in the human cohort from AUC 0.832 to 0.919.

## Introduction

Evolutionary conservation is the most widely used single predictor of whether an amino acid position matters. It decides which residues a protein engineer may move, it is a dominant feature in variant effect predictors, and it is delivered to clinical and experimental users as a per-position grade. Empirical Bayesian rate estimation on an explicit phylogeny, implemented in Rate4Site [1] and delivered at scale through ConSurf and ConSurf-DB [2, 3, 4], replaced simple counting more than two decades ago and remains the reference implementation. ConSurf marks a position computed from few ungapped sequences as unreliable and returns a confidence interval with each grade, which is the reliability statement in routine use, and it says nothing about the two failure modes characterised here. Coevolution pipelines such as EVcouplings [5, 6] report a per-column conservation value alongside their coupling statistics. The same quantity reaches the bench through general alignment tools, with Bio3D computing per-position entropy over a 22-symbol alphabet whose two extra symbols are the gap and a mask character [7], and Jalview annotating every column with a conservation value that counts a gap as a symbol carrying every property [8]. Between them these tools supply most of the conservation numbers in present use, and scoring that entropy against a background distribution instead of a uniform one is already known to change the measure [9].

What has not kept pace is the treatment of the alignment behind the number. A conservation score is computed from a sequence set somebody chose, at a depth somebody set, and in practice both are defaults. ConSurf-DB samples at most 300 homologues under a 95 per cent redundancy ceiling and a 35 per cent minimum identity, then returns grades normalised within each protein [3]. EVcouplings builds its alignments by jackhmmer against UniProt under a per-protein bitscore threshold that sets the effective depth [5]. ConSurf reports the number of sequences available at each position; the gap fraction of the column, the taxonomic composition of the set and the clade structure within it are not reported beside the score by either tool, nor in the work we have built on them [10, 11].

Neighbouring fields acted on this long ago. Alignment uncertainty propagates into genome-scale inference strongly enough to change the answer [12] and distorts sitewise tests for positive selection [13], and the response was a family of tools, Gblocks [14], trimAl, GUIDANCE and their filtering descendants, that delete unreliable and heavily gapped columns before any inference is attempted [15, 16, 17]. Conservation scoring inherited little of this hygiene. The very columns a phylogeneticist would discard as unalignable are scored, graded and reported, and a column consisting entirely of gaps is scored as maximally conserved under one widely used definition. The same gapped column is evidence too weak to place on a tree and evidence strong enough to forbid a substitution.

The two failure modes we characterise push in opposite directions (Figure 1). A column that is mostly gaps contains almost no residues, and a measure that counts the gap as a twenty-first character reads the absence of evidence as constraint, so the score comes out too high. A column whose variation is partitioned between paralogous subfamilies, fixed within each but different between them, looks variable to any measure that pools the family, so the score comes out too low as a statement about the clade the query belongs to. One mode manufactures constraint and the other conceals it. A single score can be wrong in either direction, and nothing in the output says which.

**Figure 1.**
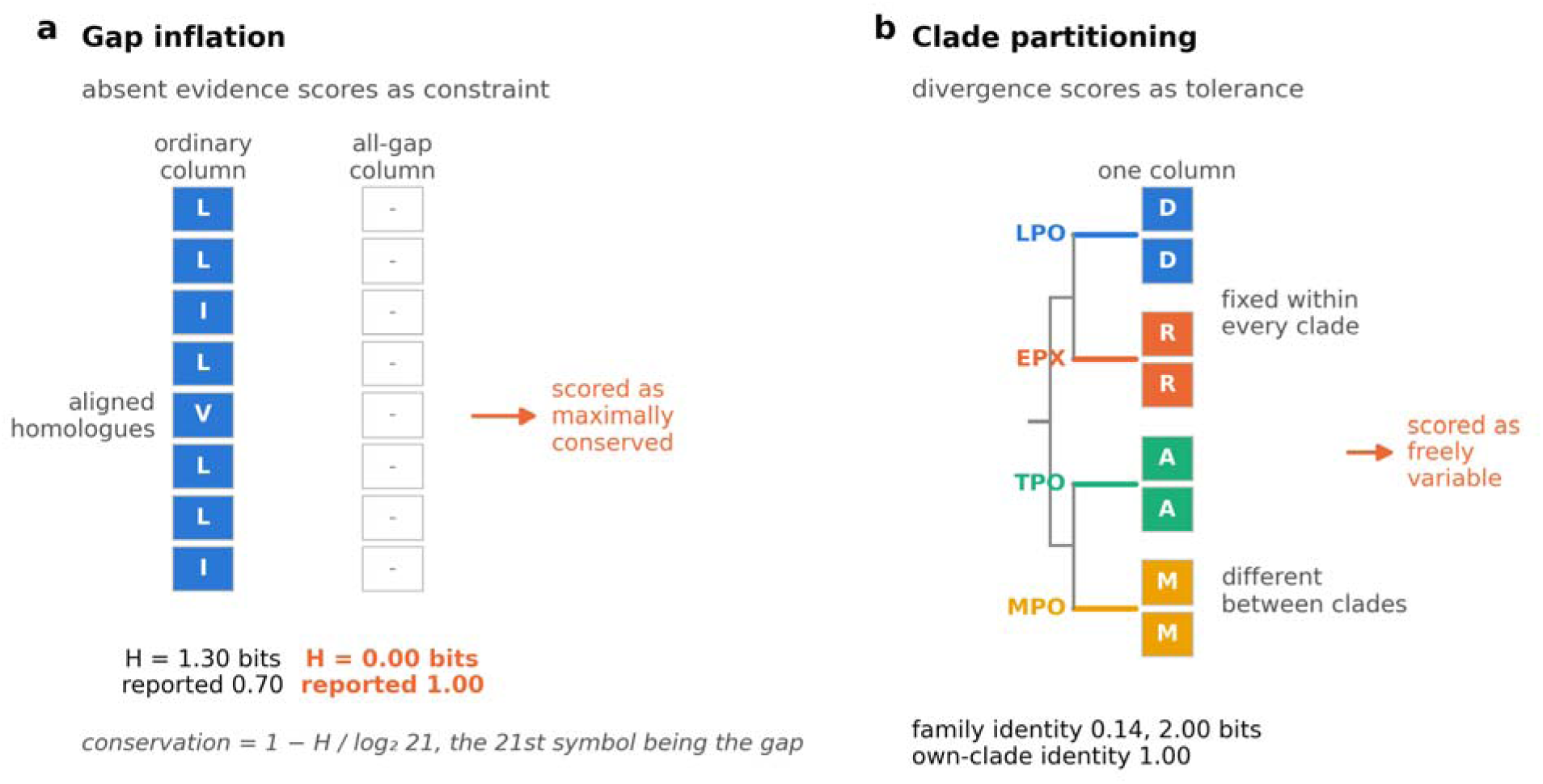
The two failure modes, and why they push in opposite directions. (a) Gap inflation. Two alignment columns over the same eight homologues. The left column holds eight ordinary residues, carries 1.30 bits of entropy and returns a conservation of 0.70. The right column holds nothing but gaps; because the gap is the twenty-first symbol of the alphabet its entropy is 0.00 bits and its conservation is the maximum attainable, 1.00. Absence of evidence is scored as constraint, and 394 of the 55,816 columns in the cohort, 0.71 per cent, are this empty. (b) Clade partitioning. One column over eight peroxidase sequences, two from each of four subfamilies, in the arrangement of lactoperoxidase D144. The residue is fixed within every clade and different between clades, so the column reports 14 per cent identity and 2.00 bits of entropy to any measure that pools the family, and is read as freely variable, while within the query’s own clade it is invariant. The real column, over sixteen comparators, gives 25 per cent identity at subfamily depth and 2.5 bits. The variation is phylogenetic, not functional tolerance.

Gap handling is not a new concern, but it has been treated as a design choice among several rather than as a measurable distortion with a sign. Valdar’s survey of conservation measures devotes a section to it and notes that the available schemes disagree [18]; Capra and Singh attached an explicit gap penalty to a Jensen-Shannon divergence score [19]; AL2CO offered several gap treatments as user options [20]. What has not been done is to establish exactly which function of the column one widely used conservation value is, measure the size and direction of the distortion that definition creates across tens of thousands of columns, and let an independent predictor arbitrate between the candidates.

The second mode has a long literature, which this work adopts instead of reinventing. Treating the partition of a family into subfamilies as informative about function goes back to the projection of sequence space onto its principal components [21] and to the evolutionary trace, which reads functional sites off the nodes of a dendrogram [22]. Hannenhalli and Russell formalised the prediction of functional subtypes from an alignment [23]; Gu placed the altered-rate-after-duplication case on a statistical footing [24]; and mutual information between residue identity at a column and a subfamily label became the statistic of SDPpred [25]. SDPhound added a permutation-based significance test [26]; SPEL scored its specificity determinants against a null simulated along a reconstructed tree, which is the nearest precedent for the null used here [27]; GroupSim reported that a simple similarity statistic matches the elaborate methods [28]; and a parallel line of work asked explicitly how functional residues might be detected independently of phylogeny [29, 30].

That literature asks which positions determine functional specificity and answers it well. It does not ask how often the phenomenon overturns a tolerance call made from a family alignment, in which direction the score is then wrong, and whether the condition can be detected before the score is used. Those are the questions here, and answering them requires a different null model.

Felsenstein’s observation that species are not independent observations, and that a comparative test must account for the tree relating them, is forty years old and routine for continuous traits [31]. It applies with equal force to a discrete character at an alignment column. A label-permutation null, which reassigns subfamily assignments across sequences, destroys all phylogenetic correlation at once and not only the part attributable to function. Label-based nulls for coevolutionary contact prediction are anti-conservative and need phylogenetically deep alignments to behave [32], and in trait-association testing simulation along the tree has displaced naive permutation for the same reason [33].

A conservation gate built from a small paralogue comparison is a routine safeguard before a residue is moved. One such gate on human lactoperoxidase, built from the three other human members of the four-gene mammalian peroxidase group, eosinophil peroxidase, thyroid peroxidase and myeloperoxidase, passes four candidate substitutions proposed by pI-code, our method for designing pH-responsive proteins, which is the subject of a patent application on which both authors are inventors [34]. An alignment of the same family read at orthologue depth reverses the identity call at all four, and the tree-aware verdicts reported below are weaker still. Position D144 is the clean case. It is aspartate in four of the five lactoperoxidases, arginine in all four eosinophil peroxidases, alanine in four of the five thyroid peroxidases and methionine and histidine in the two myeloperoxidases, with a valine and a tyrosine making up the remaining two, so it presents as freely variable at 25 per cent identity and 2.5 bits of entropy while each residue it takes is at or near fixation in the subfamily that takes it (Figure 1b). Whether that partition is more than shared ancestry is the question the rest of this paper is about, and the answer for D144 turns out to be marginal. The property that makes a position look safe to a three-paralogue test is precisely the property that makes it unsafe.

That failure generalises beyond one design, because the same alignments feed our other work and the field’s. ConSurf-DB grades entered partial correlation analyses of residue co-occurrence across 504 human alpha-helical transmembrane proteins [10], and our rigid and dynamic conserved scores multiply conservation by structural rigidity and by normalised flexibility, so an error in the conservation factor enters both [11]. We argued from a three-member human paralogue comparison to the architecture of the junctional adhesion molecules [35], and two further studies rested on pooled family alignments [36, 37]. In none of these did we state the composition of the homologue set or separate orthologues from paralogues. The same silence runs through the design literature that uses conservation as a safety gate, including the QTY substitutions we have applied to membrane proteins [38, 39]. A statistic compared against the wrong null reports an effect that is really the structure of the comparison, and the right null here costs computation and no new data.

Gap inflation is a definitional property of a widely used conservation value rather than a subtle statistical artefact, and an independent predictor arbitrates which of two candidate definitions is the distorted one. Changing only the comparator set moves a position by about two grades on the nine-bin within-protein percentile scale built as ConSurf builds its grades, with identity in place of a rate, and flips 2.7 to 8.3 per cent of positions from variable to conserved. The label-permutation null that most specificity-determining-position methods use cannot separate clade structure from shared ancestry, a tree-aware null with per-column parsimony rate matching can, and the resulting rate is governed by subfamily sampling depth and not by the family. Both modes are computable from the alignment alone, which makes them a reliability panel and not only an observation.

## Results and Discussion

### A gap-inclusive alphabet makes absent evidence read as constraint

Alignment pipelines report a per-column conservation value beside the residue frequencies they compute it from, and that value can be recovered from those frequencies. Across all 133 frequency files of our dynamics-aware cohort, 55,816 columns in total, the conservation column of the EVcouplings output is reproduced to within 0.0134 by

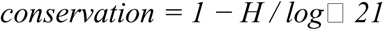

where H is the Shannon entropy of the column over a 21-symbol alphabet whose twenty-first symbol is the gap, at a minimum per-file correlation of 0.999946 and a mean of 0.999976. The residual is the rounding of the published three-decimal frequencies. The consequence is definitional and not statistical. A column that is entirely gap has H = 0 and therefore a conservation of exactly 1.000, the maximum attainable (Figure 1a).

The effect is large and monotone in the upper range. Reported conservation falls gently from 0.405 to 0.349 across the first three gap deciles, as genuinely variable positions accumulate, then climbs to 0.580, 0.776 and finally 0.934 in columns that are 90 to 100 per cent gap (Figure 2a, Table S2). Recomputed over the twenty residue symbols renormalised to the ungapped mass, the same columns stay between 0.326 and 0.418 across the first nine deciles and reach 0.597 in the most gapped one, where too few residues remain to carry entropy, which is the condition the effective-comparator criterion covers. Over the whole set the gap fraction and the reported value correlate at ρ = +0.553, falling to +0.066 after the recomputation, with both cohorts showing the same collapse independently, from +0.542 and +0.563 to −0.016 and +0.152 (Figure 2b). One fifth of all columns, 20.3 per cent, are more than half gap, and 7.61 per cent are more than half gap while reporting a conservation above 0.80.

**Figure 2.**
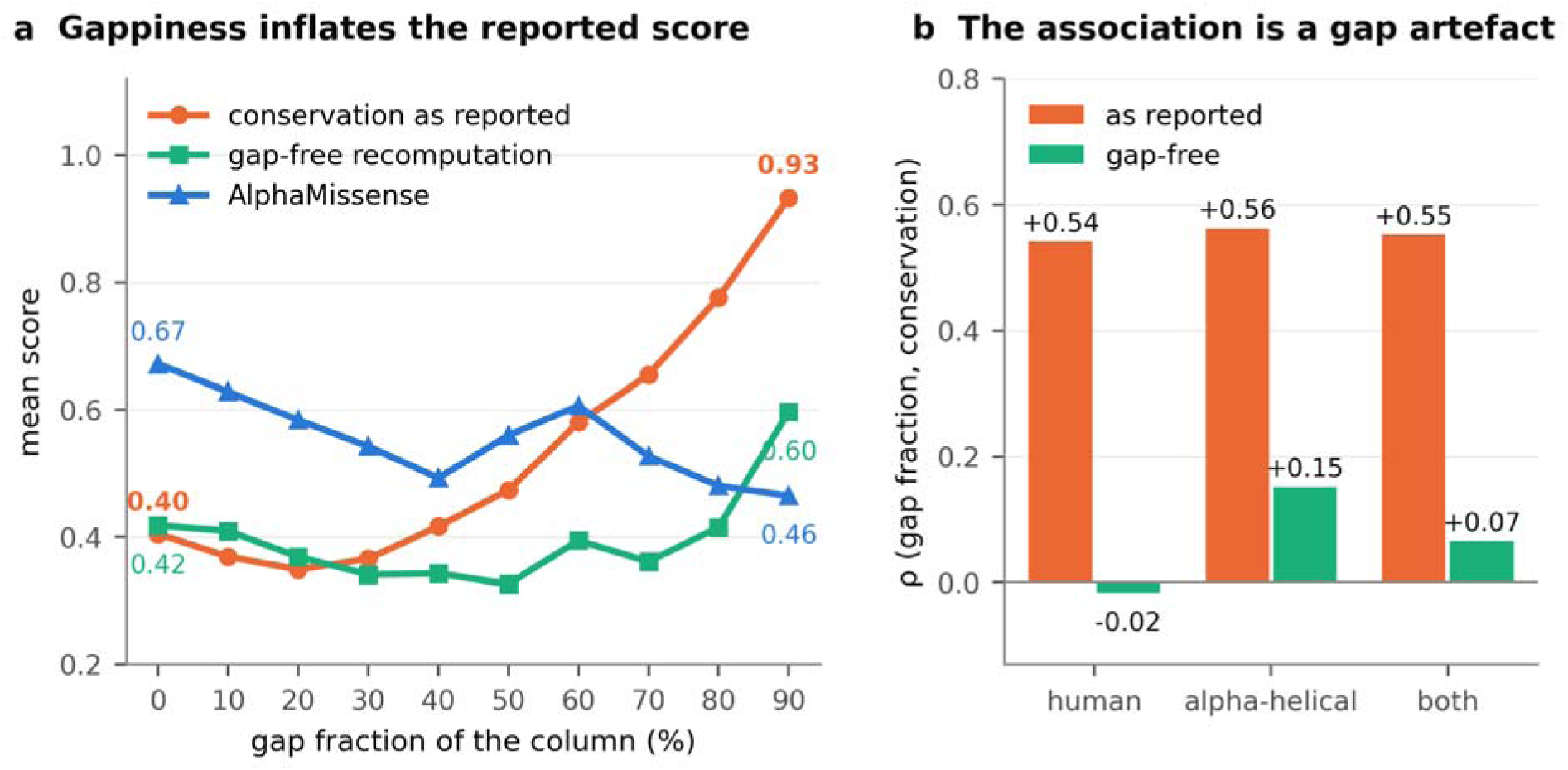
Gap inflation measured across 55,816 alignment columns. (a) Mean score by decile of column gap fraction, over the 133 EVcouplings frequency files of the dynamics-aware cohort. Conservation as reported falls gently across the first three deciles and then climbs steeply, reaching 0.934 in columns that are 90 to 100 per cent gap. The gap-free recomputation, which renormalises the twenty residue frequencies to the ungapped mass, stays between 0.326 and 0.418 across the first nine deciles and reaches 0.597 in the most gapped one where too few residues remain to carry entropy. AlphaMissense, an independent judgement, falls from 0.671 in the least gapped decile to 0.465 in the most, against a reported conservation that climbs to 0.934 over the same range. (b) The correlation between column gap fraction and conservation, before and after the recomputation, in the human and alpha-helical cohorts and in the pooled set. The association is a definitional artefact of the gap-inclusive alphabet, not a property of the proteins.

Alignment depth is the obvious candidate for a common cause, on the reasoning that a permissive search admits distant sequences covering less of the query, and it was registered as a prediction before the test. The per-protein jackhmmer bitscore threshold varies sevenfold across this cohort, from 0.1 to 0.7, and is not reported in the source publication. Its rank correlation with per-protein mean gap fraction is −0.034 over 133 proteins (Figure S3), so the search threshold does not predict gap inflation, and the gap mode is not a by-product of the depth parameter. A panel therefore has to report the gap fraction itself, since the search threshold the alignment was built at does not predict it.

### An independent predictor arbitrates between the two definitions

Whether a predictor inherits the gap flaw is a separate question from whether the conservation value has it, and we registered both branches before computing: a positive association between gap fraction and AlphaMissense score would indicate that the predictor shares the flaw, a negative one that the predictor handles gaps correctly while the conservation value does not [40]. Over 13,051 residues in 70 proteins the association is negative, ρ = −0.223, and remains negative at −0.170 after partialling out the gap-free conservation. AlphaMissense treats gappy positions as less constrained, the direction a gap-free measure predicts.

The discriminating comparison is which conservation definition better tracks that independent judgement. The reported value reaches ρ = 0.416 and the gap-free recomputation ρ = 0.551, a gain of 0.135 that appears independently in both cohorts, 0.435 to 0.571 in the human set and 0.431 to 0.558 in the alpha-helical one (Table 2). Removing the gap from the alphabet brings the measure substantially closer to AlphaMissense, which identifies the reported value as the distorted one without any appeal to the mechanism. In 121 curated human variants mapped to these positions, with every wild-type residue agreeing with its sequence, AlphaMissense scores lower at gappy positions in both classifications, 0.297 against 0.437 among the 49 benign-side variants and 0.575 against 0.793 among the 52 pathogenic-side ones, consistent in direction at low power. The remaining 20 are of uncertain significance and carry no classification to compare.

**Table 1.** Peroxidase positive control and the four designed positions under the tree-aware null.

| Position | Set<sup>1</sup> | H bits<sup>2</sup> | sef<sup>3</sup> | Null mean<sup>4</sup> | Corrected<sup>5</sup> | p<sup>6</sup> | Call |
| --- | --- | --- | --- | --- | --- | --- | --- |
| Ten catalytic and cofactor positions | control | 0.000 | - | - | - | - | conserved |
| D144 | walked | 2.500 | 0.770 | 0.521 | 0.518 | 0.027 | diagnostic, marginal |
| E648 | walked | 0.811 | 1.000 | 0.378 | 1.000 | 0.134 | indeterminate |
| E205 | walked | 0.868 | 0.766 | 0.381 | 0.623 | 0.046 | diagnostic, marginal |
| E313 | walked | 0.896 | 1.000 | 0.353 | 1.000 | 0.150 | indeterminate |
| D479 | held back | 0.696 | 0.385 | 0.384 | 0.001 | 0.474 | tolerant |
| E169 | held back | 0.000 | - | - | - | - | conserved |
| D700 | variant | 1.717 | 0.619 | 0.507 | 0.228 | 0.248 | tolerant |

<sup>1</sup>Set names the role of the position in the original design: control, one of the four walked positions, one held back from the design, or a known human variant position.
<sup>2</sup>H is the entropy of the column in bits.
<sup>3</sup>sef is the subfamily-explained fraction, the mutual information between residue identity and subfamily label normalised by the column entropy.
<sup>4</sup>Null mean is the mean subfamily-explained fraction of the simulated neutral character over 1,000 replicates at the calibrated rate.
<sup>5</sup>Corrected is the chance-corrected value, (sef – null mean) divided by (1 – null mean), floored at zero.
<sup>6</sup>p is the empirical significance of the observed value against the tree-aware null. A dash means the column is below the entropy floor of 0.5 bits and carries no value.

**Table 2.** Rank correlation of two conservation definitions with AlphaMissense, by cohort.

| **Cohort<sup>1</sup>** | **Residues** | **$\rho(\text{reported, AM})^2$** | **$\rho(\text{gap-free, AM})^3$** | **$\rho(\text{gap, AM})$** | **Partial $\rho(\text{gap, AM})^4$** |
| --- | --- | --- | --- | --- | --- |
| Human | 10,213 | 0.435 | 0.571 | -0.172 | -0.115 |
| Alpha-helical | 2,838 | 0.431 | 0.558 | -0.361 | -0.253 |
| Both | 13,051 | 0.416 | 0.551 | -0.223 | -0.170 |

<sup>1</sup>Human is the 504-protein human set and alpha-helical the transmembrane set, both as assembled in our previous work [10, 11]; Both is the two pooled, 70 proteins in all.
<sup>2</sup>Reported is the conservation column as the alignment pipeline supplies it, $1 - H/\log 21$ over an alphabet whose twenty-first symbol is the gap.
<sup>3</sup>Gap-free is the same column recomputed over the twenty residue symbols renormalised to the ungapped mass, $1 - H/\log 20$ .
<sup>4</sup>Partial is the correlation between gap fraction and AlphaMissense with gap-free conservation partialled out. AM is AlphaMissense [40]. All correlations are Pearson on ranks, and partial correlations the rank-based partial form.

AlphaMissense is itself trained on multiple sequence alignments, and a predictor validated against the alignments it learned from can confirm its own artefacts; the two documented forms of circularity in variant-effect benchmarking are exactly of this kind [41]. The argument survives because the direction is wrong for circularity. If AlphaMissense had absorbed the gap-as-conservation convention it would score gappy columns higher, not lower, and the reported value would track it better than the gap-free one. It does the opposite in both cohorts. Deep generative models of evolutionary data behave the same way for the same reason, since a gapped column supplies them with no residue to model [42]. Gap inflation is therefore a defect of conservation measurement and of the scores derived from it, and not of the predictor.

### The gap artefact propagates into published derived scores

Because the rigid and dynamic conserved scores are defined as the product of conservation with rigidity and with normalised flexibility [11], the distortion enters both multiplicatively. Taking that paper’s own published score table as the population, and counting the residues on columns more than half gap: against a base rate of 15.7 per cent in the human cohort, the top decile of the dynamic conserved score is 44.7 per cent such residues and its top percentile 70.9 per cent; in the alpha-helical cohort, against a base of 14.4 per cent, the same tails are 49.3 and 79.2 per cent (Figure S4b). The rigid score is enriched in the same direction, at 40.9 and 45.0 per cent in the top decile of the two cohorts. Roughly half the residues in that paper’s top decile, and three quarters of those in its top percentile, therefore sit on columns that are more than half gap, in both cohorts independently.

We rebuilt that pipeline four ways, renormalising per protein exactly as published: as published, with gappy columns filtered, with gap-free conservation, and with both (Table S7). The as-published variant reproduces the published independence of the two scores at ρ = 0.080 against the reported 0.08, and their correlation with raw conservation at 0.903 against 0.91. The reimplementation therefore reproduces the published pipeline. Handling the gap then improves the published score’s own discrimination, with the rigid conserved score moving from AUC 0.832 (95 per cent confidence interval 0.816 to 0.848) to 0.919 (0.909 to 0.928) in the human cohort and from 0.770 to 0.928 in the alpha-helical one, while top-percentile precision for the dynamic score rises from 0.406 to 0.778 (Figure S4a). The dynamic score’s own area under the curve remains at chance against this label, 0.438 to 0.541 across the four variants in the human cohort and 0.487 to 0.569 in the alpha-helical one, which is what that paper itself reports. Table S9 gives the same four variants against the clinical label.

Removing the gap from the alphabet therefore raises the discrimination available to that pipeline and clears its top ranks of columns carrying almost no residue. Two of the four variants named in its discovery table, NARS1 p.Thr17Met and p.Lys60Thr, sit on columns that are 98.5 and 93.4 per cent gap, leaving the residue present in 1.3 and 6.5 per cent of the aligned mass, which is the case the panel’s effective-comparator criterion covers and where no estimate is available in either direction. A 50 per cent gap filter excludes both columns from the analysis instead of scoring them low. The other two, PGK1 p.Lys30Thr and SPRTN p.Tyr117Cys, at 4.9 and 0.1 per cent gap, are unaffected. A derived score inherits the reliability of its inputs.

Neither measurement rests on a hypothetical implementation. The definition was read off the published frequency output of the EVcouplings pipeline across 55,816 columns, and the derived scores are the ones the source paper reports, so what is at issue is not whether a gap-inclusive alphabet is advisable but what it has already done to published results.

### Changing only the comparator set moves the conservation call

To isolate the second failure mode we held the reference, the aligner and the guide-tree method constant, remapped positions to the reference row independently at each depth, and varied only which sequences were compared, drawing all of them from one source file so that the sequences themselves were identical between depths. Over human lactoperoxidase this gives four comparator sets, the first two disjoint and both contained in the third: the three other human peroxidases, which is the gate the original design ran under; the five non-human lactoperoxidases; the sixteen other mammalian-type peroxidases; and the seventy-nine other reviewed members of the family. 558 of 712 positions pass the gap filter at every depth.

The structure of the movement is what the statistic predicts (Figure 3a). Positions the subfamily-explained fraction calls diagnostic invert sharply between the paralogue gate and orthologue depth, from a mean identity of 0.256 to 0.815, while positions it calls tolerant stay nearly flat, 0.797 to 0.757. The measure separates the positions the change in depth affects from those it does not. Against orthologue depth, 22.4 per cent of positions invert at the three-paralogue gate, 17.4 per cent at subfamily depth and 49.1 per cent across the full family, so the rate is not monotone in depth and the widest alignment is the worst. Positions invariant across all seventeen mammalian-type peroxidases fall to a mean identity of 0.527 in the eighty-sequence alignment.

**Figure 3.**
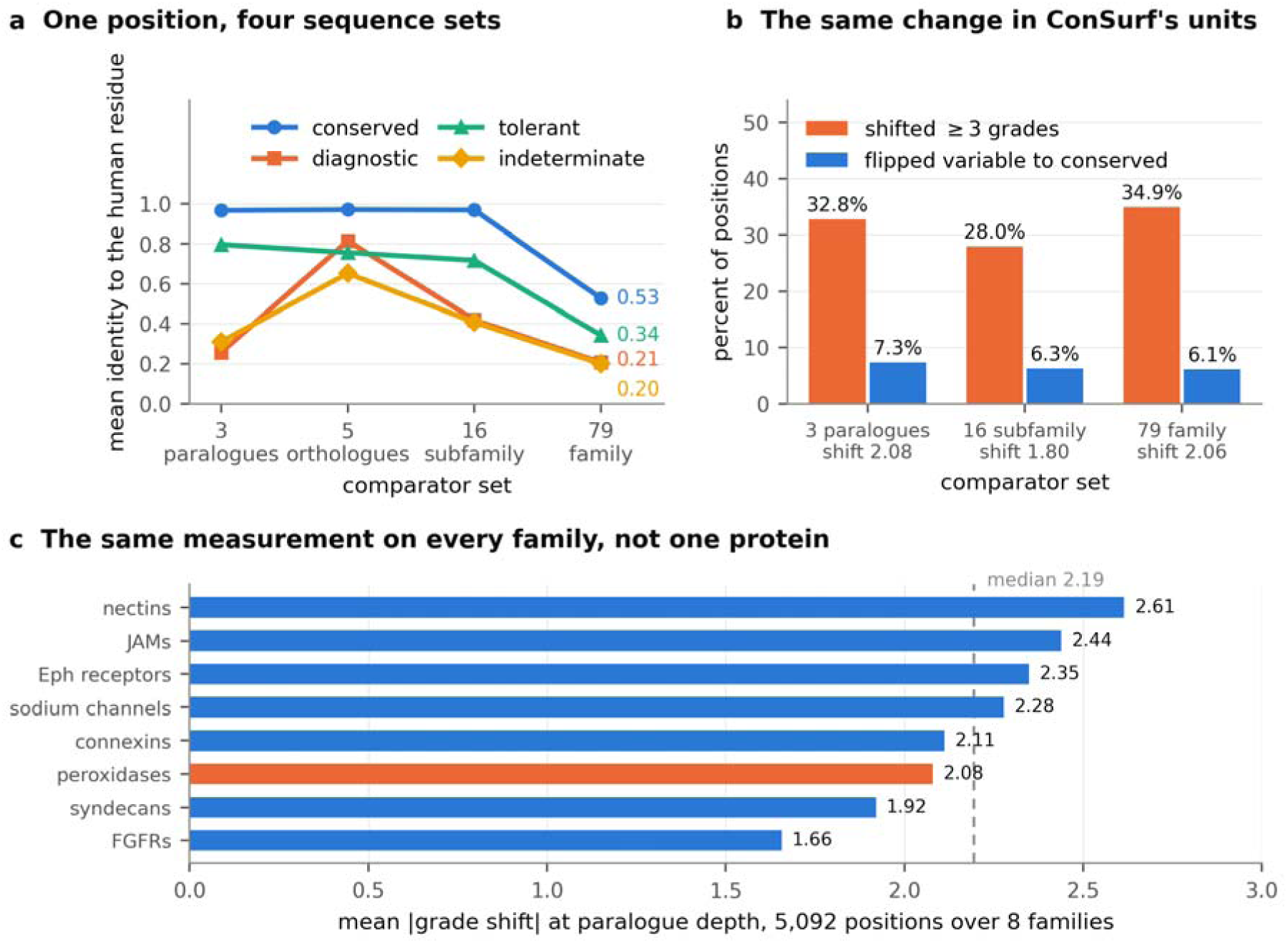
Changing only the comparator set moves the conservation call. (a) Mean identity of the human lactoperoxidase residue to the comparison set, at four nested depths, with positions grouped by the call the subfamily-explained fraction makes at orthologue depth. Positions called diagnostic invert sharply between the three-paralogue gate and orthologue depth, from 0.256 to 0.815, while positions called tolerant stay nearly flat. The statistic separates exactly the positions that the change in depth affects. (b) The same comparison expressed in ConSurf’s own units. Grades are equal-count percentile ranks within a protein and are therefore scale-free, so a disagreement between depths is a disagreement about the ordering of positions and not about the divergence of the set. Changing only the comparator set shifts 28.0 to 34.9 per cent of positions by three grades or more and flips 6.1 to 7.3 per cent the whole way from variable to conserved. (c) The same measurement repeated on every family rather than on one protein. Mean absolute grade shift between the paralogue gate and the subfamily set, over 5,092 positions in eight families, with lactoperoxidase marked. The median family moves 2.19 grades, the median fraction moving three grades or more is 36.6 per cent, and the median flip from variable to conserved is 6.6 per cent. Lactoperoxidase is not the extreme family on any of the three, so the sensitivity is a property of the measurement and not of this protein.

Raw identity invites the objection that a more divergent set must show lower identity, so we repeated the comparison in the nine-bin within-protein normalisation ConSurf applies, with identity in place of a Rate4Site rate. Grades 1 to 9 are percentile ranks of conservation within one protein, hence scale-free, and a disagreement between grades computed at two depths is a disagreement about the ordering of positions rather than about the divergence of the set. Changing only the comparator set moves a position by a mean of 1.80 to 2.08 grades, moves 28.0 to 34.9 per cent of positions by three grades or more, and flips 6.1 to 7.3 per cent the whole way from variable, grade 3 or below, to conserved, grade 7 or above (Figure 3b). The flip rate is stable across all three comparator sets. The subfamily-explained fraction predicts the size of the movement and not only its presence, at ρ = 0.406, 0.430 and 0.353 against the identity drop from orthologue depth, all with p below 10 ¹², over 386 variable positions.

One protein cannot establish that two grades is typical, so we repeated the measurement on each of the eight assembled protein families, every one against its own paralogue depth, meaning the other human members of that family, one per gene, which is the comparator set a design has when nothing deeper is to hand. Over 5,092 positions in eight families the mean absolute grade shift runs from 1.66 in the fibroblast growth factor receptors to 2.61 in the nectins, with a median of 2.19, and the share moving three grades or more runs from 23.8 to 44.3 per cent with a median of 36.6 per cent (Figure 3c). Lactoperoxidase sits at 2.08 and 32.8 per cent, mid-range on both, so the protein the rest of this paper works on is not a peculiar one. The flip from variable to conserved runs from 2.7 to 8.3 per cent with a median of 6.6. Running the generic implementation on the peroxidases reproduces every number the detailed one reports, so generalising the measurement leaves it unchanged.

Mapped onto the structure, the flagged positions are not a localised patch (Figure 4). Clade-partitioned positions are distributed over the whole lactoperoxidase fold, including the face that carries the catalytic machinery, and the gap-inflated ones concentrate in the terminal and loop regions where the family’s members differ in length (Figure 4b). A user inspecting a conservation colouring of this structure would see the two artefacts as ordinary signal.

**Figure 4.**
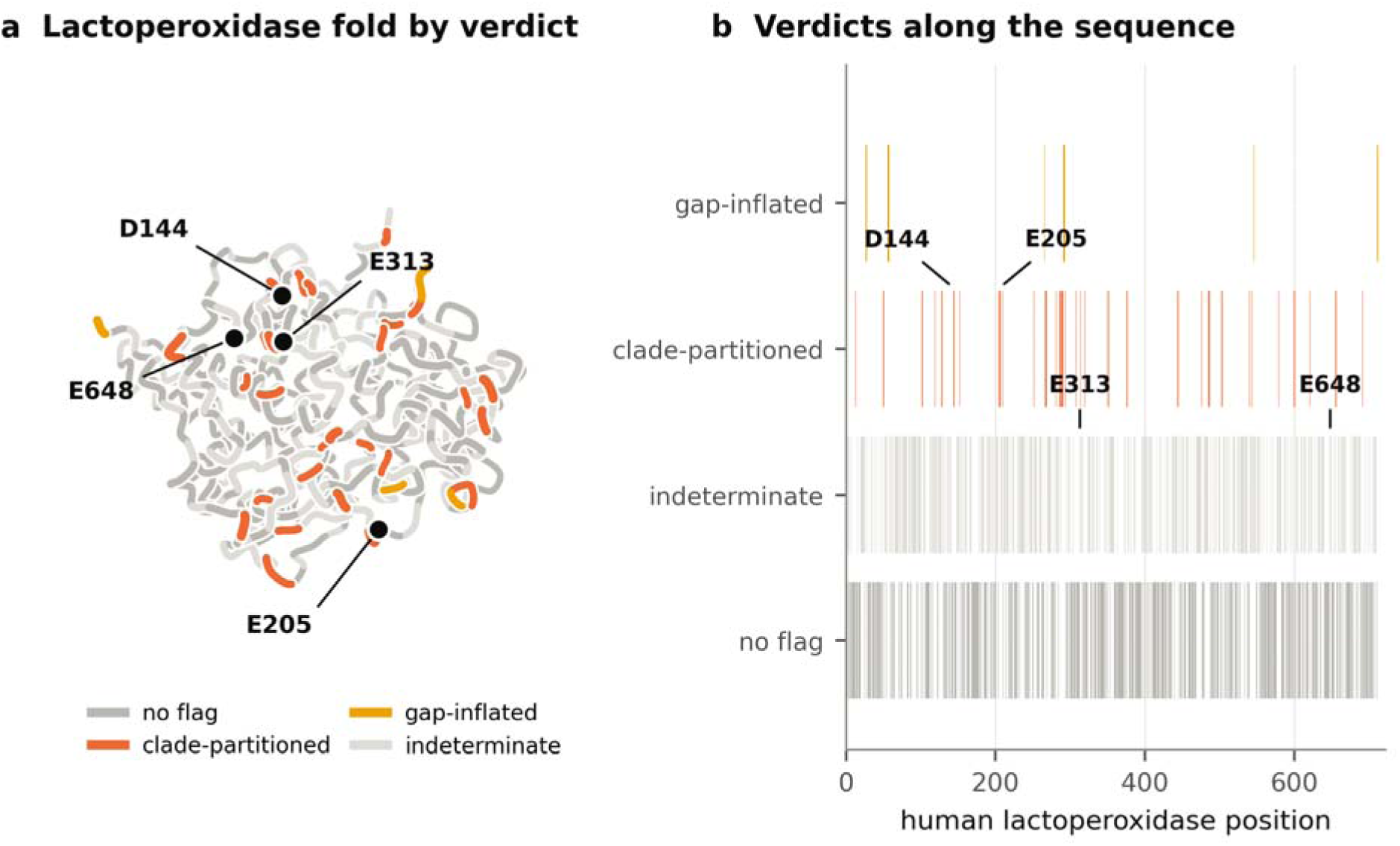
Reliability verdicts mapped onto the lactoperoxidase fold and sequence. (a) The alpha-carbon backbone of lactoperoxidase, PDB 7DE5 chain A mapped to human numbering, projected onto its two leading principal axes and drawn in depth order. Each segment is coloured by the verdict the reliability panel returns for that position: no flag, clade-partitioned under the tree-aware null, gap-inflated, or indeterminate. The four positions carried through the original design are marked. Flagged positions are distributed across the whole fold rather than confined to a surface patch. (b) The same verdicts along the sequence, one lane per verdict, with the four marked positions annotated. Gap-inflated columns concentrate in the terminal and loop regions where family members differ in length; clade-partitioned columns do not.

### A label-permutation null cannot separate clade structure from ancestry

The standard significance test for this statistic permutes subfamily labels across sequences [26]. That null asks whether these residues are more clade-partitioned than a random assignment of them to these clades, and it destroys all phylogenetic correlation at once and not only the part attributable to function. Any character evolving along a gene tree is partly clade-partitioned through shared ancestry alone [31], so the bar is set far too low. On the peroxidase family it calls 68.3 per cent of variable columns diagnostic; across nine family runs the figure ranges from 68.3 to 100 per cent (Figure 5b, Table S6). The permutation null’s expectation sits below the observed value at almost every column (Figure S2). A flag that fires on two thirds to all of the variable positions is not a flag. The tree-aware null meets that same test in one family, returning 69.1 per cent in the fourteen-subfamily Eph receptors, so the panel is informative only where the decomposition is coarse enough to leave most columns unexplained.

**Figure 5.**
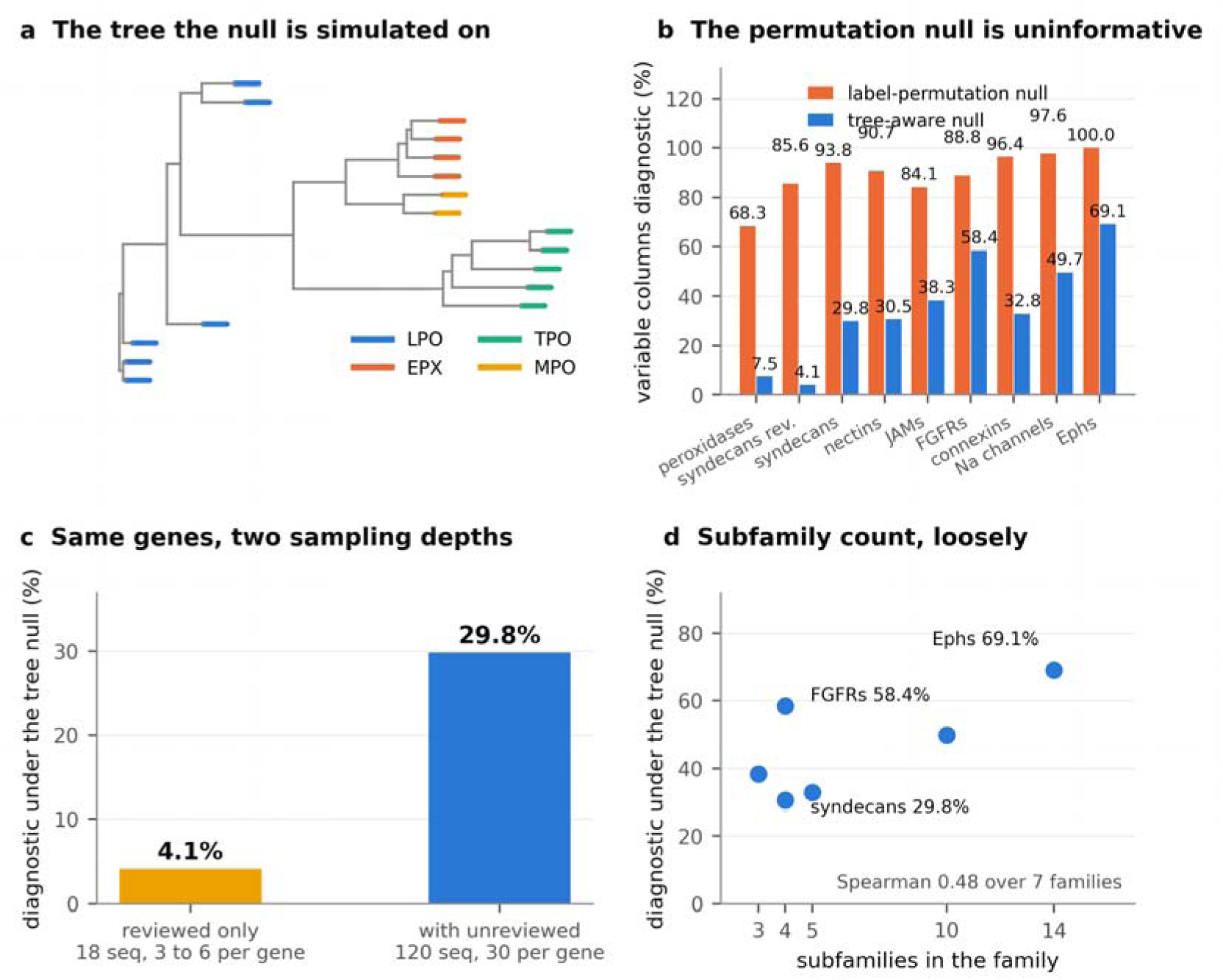
The null model determines the answer, and sampling determines the rate. (a) The neighbour-joining tree of the seventeen mammalian-type peroxidases on which the tree-aware null is simulated, with tips coloured by subfamily. The label-permutation null destroys clade structure of this kind. (b) Percentage of variable columns called diagnostic under each null, across nine family runs. The permutation null returns 68.3 to 100 per cent, which is not a usable flag; the tree-aware null returns 4.1 to 69.1 per cent. (c) The same four syndecan genes, the same reference, thresholds and code, differing only in the orthologue set: 4.1 per cent at 18 sequences and 29.8 per cent at 120. The shallow figures are floors, not estimates. (d) Tree-null rate against the number of subfamilies the family is decomposed into, over the deeply sampled families. The Spearman is 0.48, which seven families cannot resolve: the fibroblast growth factor receptors return 58.4 per cent from four subfamilies where the nectins and syndecans return 30.5 and 29.8 per cent from four. No single proteome-wide rate is well posed without stating the decomposition.

We therefore simulate a neutral character along each family’s own tree (Figure 5a). The tree is built by neighbour joining on Poisson-corrected p-distances [43], the substitution model is an F81 process whose equilibrium frequencies are the column’s own composition [44], and branch lengths are scaled per column so that the simulated character carries the same Fitch parsimony score as the observed one [45]. Conditioning the equilibrium on the column’s own composition is deliberate, because the question concerns the arrangement of these residues among clades and not which residues they are, so holding composition fixed isolates clade partitioning from the composition confound. Because F81 is reversible and stationary, the simulated tip distribution does not depend on where the tree is rooted, so the unrooted neighbour-joining output needs no root to be chosen.

The calibration quantity is not a free choice, and getting it wrong inverts the conclusion silently. Matching the simulated and observed column entropies is the obvious choice and cannot work. With the equilibrium set to the column’s own composition, the tips at high rate become independent draws from that composition, and the mean plug-in entropy of sixteen such draws lies below the plug-in entropy of the observed sixteen by the finite-sample bias, approximately (k − 1)/(2n ln 2) bits, about 0.27 bits for a seven-state column. The target is unreachable, the search saturates at the top of the rate grid, and at the top of the grid no phylogenetic correlation survives, so the tree null silently becomes the permutation null. Every column whose target cannot be reached is reported as a calibration failure, which is the only visible symptom. Parsimony rises monotonically from zero and is reachable. Across the nine runs it leaves 285 of 4,183 attempted columns uncalibrated, 6.8 per cent, ranging from 0.6 per cent on the peroxidases, 3 of 495, to 15.1 per cent on the Eph receptors, 129 of 854, and 18 columns in total settle at an edge of the rate grid (Figure S1a). An analysis calibrated on entropy instead returns the permutation null under a different name, and Note S1 gives the argument in full.

Clade partitioning is not selection even under a tree-aware null. The null establishes that a column is more clade-structured than a neutral character on that tree, which is a statement about measurement and not about function, and the word diagnostic should be read throughout as clade-partitioned.

### Sampling depth moves the clade-partitioning rate as much as family identity

Under the tree-aware null the peroxidase rate falls from 68.3 to 7.5 per cent of variable columns, and none of the four walked positions survives as unambiguously diagnostic. D144 and E205 sit at chance-corrected values of 0.518 and 0.623 with p of 0.027 and 0.046, marginally past the declared threshold. An estimate sitting this close to its own threshold is at the limit of what seventeen sequences can resolve. E648 and E313 go the other way. Both partition perfectly, with a subfamily-explained fraction of exactly 1.000, and both are unremarkable against the tree null at p = 0.134 and 0.150, because with four clades and few substitutions a neutral character reaches a perfect partition often (Table 1).

At the reviewed tier NECTIN2 and JAM2 retain two mammalian entries each, which meets the floor of two ungapped sequences only at columns where neither is gapped and leaves the within-group entropy resting on a single pair. Across nine runs covering eight families the permutation null calls 68.3 to 100 per cent of variable columns diagnostic while the tree-aware null calls 4.1 to 69.1 per cent (Table S6, Figure 5b). The three families added to test the sister-state prediction, the connexins, the sodium channel alpha subunits and the fibroblast growth factor receptors, fall at 32.8, 49.7 and 58.4 per cent, inside that range and not at either end.

Building the families makes the orthology criterion part of the result, so it is validated against a stricter criterion instead of being taken on trust. Ensembl Compara calls orthology from reconciled gene trees [46, 47], and of the 1,102 non-human sequences in the eight families it can adjudicate 765, confirming 757 of them as one-to-one orthologues, 99.0 per cent, with per-family rates from 98.3 to 100 per cent. The eight exceptions are orthologues that are not one-to-one, in six species, goat and kangaroo rat twice each and wombat, rat, chimpanzee and brown bear once, which is the lineage-specific duplication a gene-name criterion admits by design. Of the 337 it cannot adjudicate, 307 are in species for which it holds no genome and 30 are non-mammalian members of the Pfam-derived peroxidase file. Note S2 sets the criterion out in full, Table S1 gives what each family assembly kept and rejected, and every rejected sequence is listed with its reason in Supplementary Table S18.

The decisive comparison holds the family fixed. The same four syndecan genes, the same reference, the same thresholds and the same code, differing only in the orthologue set, give 4.1 per cent at 18 sequences and 29.8 per cent at 120 (Figure 5c). The deep families rest on 10 to 18 curated entries out of 90 to 280, so curation sets the floor and the unreviewed tier sets the ceiling. A sevenfold swing from sampling depth alone in the one family measured at two depths means the shallow numbers, including the peroxidase 7.5 per cent, should be read as lower bounds until a second family is measured the same way. It also means that no single proteome-wide rate is well posed without conditioning on how the family was decomposed and how deeply each subfamily was sampled. Subfamily count is part of it, and the fourteen-subfamily Eph family does return the highest rate at 69.1 per cent. Over the seven deeply sampled families the association between subfamily count and the tree-null rate is a Spearman of 0.48, which seven families cannot resolve, and the clearest exception is the fibroblast growth factor receptors, which return 58.4 per cent from four subfamilies where the nectins and syndecans return 30.5 and 29.8 per cent from four (Figure 5d). How far the paralogues have diverged is the obvious candidate alongside how many of them there are.

Detecting clade partitioning requires subfamily sampling that shallow alignments do not have, and the scores most often used are computed from exactly those alignments. ConSurf-DB caps at 300 homologues with no subfamily structure at all [3], and the EVcouplings bitscore that sets depth varies sevenfold across a single published cohort without being reported [11].

### A per-position reliability panel, and a directional prediction

The panel reports three quantities for every scored position. The gap fraction of the column, whose exceedance means the score is too high. The effective comparator count after gaps are removed, whose shortfall means there is no estimate in either direction. The tree-corrected subfamily-explained fraction with its empirical p, whose exceedance means the score is too low as a statement about the query’s own clade. The first two are already available wherever a conservation score is computed, and the third requires a tree and a per-column simulation. On the peroxidase family the panel flags 5.2 per cent of positions as clade-partitioned and 1.3 per cent as gap-inflated, calls 15.0 per cent tolerant and 29.6 per cent conserved with no subfamily-explained fraction defined, and leaves 48.9 per cent indeterminate under the tree-aware null. At this sampling depth the largest class is the one the panel declines to call. The five-comparator floor cannot fire on a sixteen-sequence set once the gap threshold is applied, since the assessed columns carry 12 to 16 comparators, so that criterion is untested here and matters at the alignment depths most scores are computed from. Scored with the label-permutation null in place of the tree-aware one the same panel flags 336 of the 712 positions, 47.2 per cent, as clade-partitioned, and 35 of the 37 the tree-aware null flags lie among them, so the permutation null flags roughly nine times as many positions as survive it. Figure 4a carries the tree-aware verdicts on the lactoperoxidase fold, with the positions the permutation null alone flags shown as indeterminate.

The panel also emits an annotation that generates a testable biological prediction, and with eight families assembled there are now enough curated variants to test it. At a clade-partitioned position, each sister subfamily fixed at a residue unlike the query’s supplies a substitution the fold demonstrably accepts somewhere in the family; 390 of 712 peroxidase positions carry at least one such sister state, 34 of them clade-partitioned under the tree-aware null and 307 under the permutation null. Lactoperoxidase E648 is the worked example of the annotation, at a position the tree-aware null leaves indeterminate (Table 1): glutamine is the fixed state of all four eosinophil peroxidases, and the sister state exists whatever the null says about the column. The prediction, registered in the analysis script before the pooled numbers existed, is that at a clade-partitioned position human missense variants to a sister subfamily’s fixed state are enriched for benign classification relative to other substitutions at the same positions.

The ten most frequent ordered sister substitutions across the eight families are V→I, I→V, T→S, A→S, I→L, D→E, E→D, K→R, S→A and V→L, every one a conservative exchange and together 16.8 per cent of the 8,649 instances at the 3,063 positions carrying a sister state (Table S3), so a pooled test would partly rediscover that conservative substitutions are benign. Ascertainment is a second confound. A disease gene is sequenced in patients, so the benign share among its curated variants runs from 0.084 in the connexins to 0.762 in the Eph receptors (Table S8, Figure 6a), and pooling those into one table would compare families rather than substitutions. Both are handled by stratifying on family and on substitution conservatism and combining the strata by Mantel-Haenszel, which conditions on each stratum’s own margins; Note S3 sets out what that costs and why both arms are reported.

**Figure 6.**
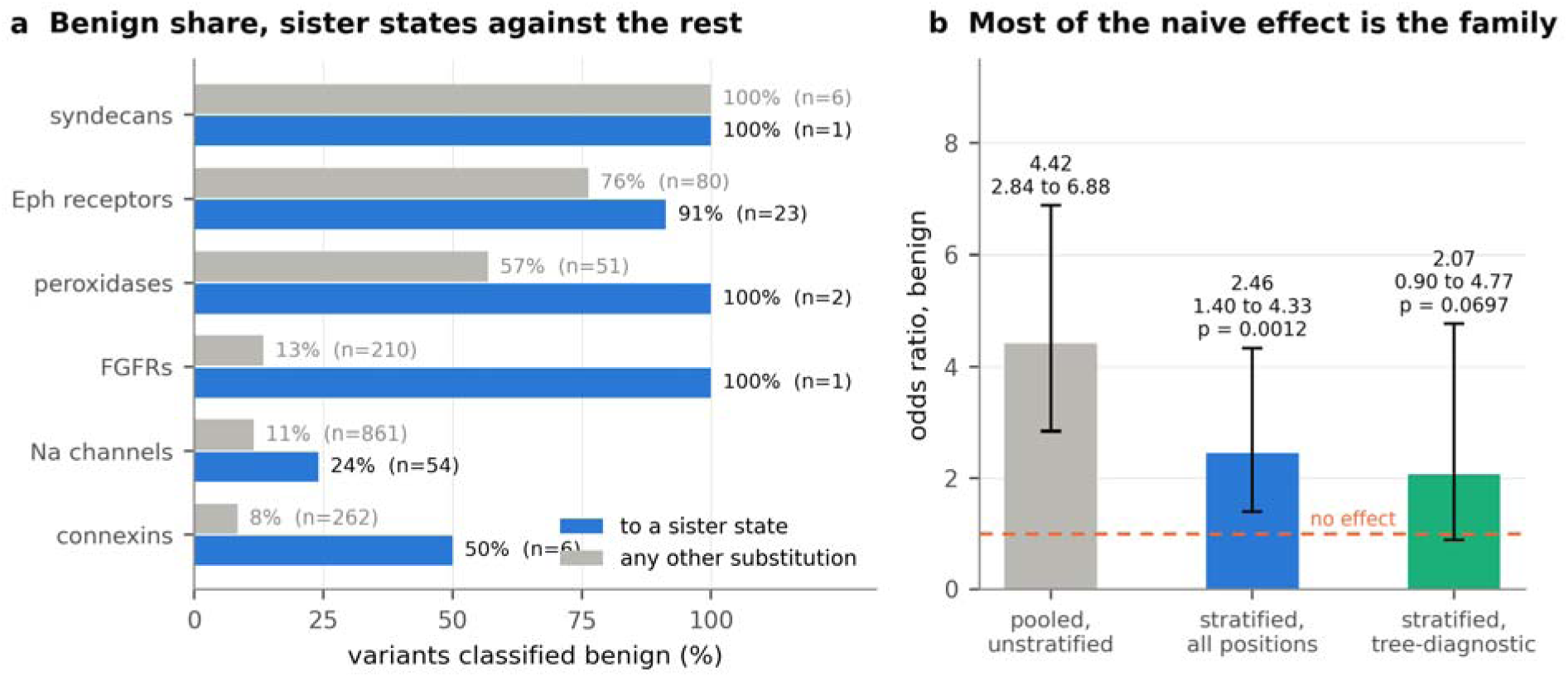
Variants to a sister subfamily’s fixed state are enriched for benign classification. (a) Percentage of curated human missense variants classified benign, sister-state substitutions against every other substitution, by family, with the number of classified variants behind each bar. The benign base rate differs ninefold between families because a disease gene is sequenced in patients, which is why the families are compared only against themselves. The syndecan, peroxidase and FGFR rows rest on one or two sister-state variants and carry almost no weight in the estimate. (b) The pooled odds ratio before and after that stratification, and under the stricter reading, each with its 95 per cent interval. Unstratified, 4.42 (2.84 to 6.88) on a Woolf interval Stratified on family and substitution conservatism, 2.46 (1.40 to 4.33) on a Robins-Breslow-Greenland interval, at a one-sided p of 0.0012. Restricted further to positions the tree-aware nul calls diagnostic, 2.07 (0.90 to 4.77) at a one-sided p of 0.070, an interval that includes no effect. More than half the naive effect is the difference between families and not the substitution.

Stratification changes the estimate by enough to decide how it is read. Pooled and unstratified, the odds ratio is 4.42; stratified, it is 2.46 (Figure 6b). More than half the naive effect was the ascertainment difference between families and not the substitution. The stratified estimate over 1,565 classified mapped variants in eight families, six of which carry a sister-state variant at all, is an odds ratio of 2.46 with a one-sided p of 0.0012 and a 95 per cent Robins-Breslow-Greenland interval of 1.40 to 4.33, over nine of sixteen family-by-conservatism strata, which Table S4 gives in full; the seven remaining strata carried an empty margin and contributed nothing. Every family with more than one sister-state variant points the same way, across benign base rates differing by a factor of nine.

That denominator is the classified variants at the assessed positions, and the exposed arm within it is smaller: 87 variants are substitutions to a sister state, and 77 of those sit in two families, the Eph receptors and the sodium channel alpha subunits. The estimate is therefore carried by two families, and two restrictions test what that costs. Dropping the Eph receptors entirely leaves 2.42 on the remaining seven. Keeping only strata holding at least five sister-state variants, which removes the families resting on one or two, gives 1.94 with an interval of 1.05 to 3.60. The direction survives both and the magnitude falls, so the pooled 2.46 is the upper end of what these data support.

Restricting to positions the tree-aware null calls diagnostic, the restricted form in which the prediction was registered, leaves the direction unchanged at an odds ratio of 2.07, interval 0.90 to 4.77, and costs the significance, one-sided p 0.070 on 320 variants. We report both and treat the unrestricted arm as primary, because the mechanism is the sister state itself, a residue the fold is observed to accept in a related clade, and that property exists at a position whatever the null later says about it.

The AUC of 0.832 against 0.919 is measured against a label taken from AlphaMissense, and AlphaMissense is also the predictor that arbitrates between the two conservation definitions earlier in this work. That figure is agreement with AlphaMissense and carries no claim of clinical accuracy. Measured instead against ClinVar germline classifications [48], which are independent of both, the rigid conserved score reaches an AUC of 0.535 (95 per cent confidence interval 0.403 to 0.665) as published and 0.610 (0.482 to 0.732) once conservation is recomputed gap-free, over the 86 residues of the human cohort that carry an unambiguous classification, 34 pathogenic and 52 benign. The gap correction moves the score in the same direction against a clinical label as against AlphaMissense. The absolute discrimination is close to chance and the two marginal intervals overlap, but both are measured on the same 86 residues, so the quantity with an interpretable interval is the difference between them. Resampling those residues and recomputing both scores on each resample gives a gain of 0.075 with a 95 per cent interval of 0.003 to 0.149, which excludes zero. The clinical gain is therefore supported and its lower bound sits close enough to zero that 86 residues bound its size only loosely. Table S9 reports all four variants and both scores against this label. The dynamic score, which stays at chance against the AlphaMissense label, reaches 0.639 (0.514 to 0.759) there once conservation is recomputed gap-free, so that score’s failure to discriminate is a property of the label it was measured against and not of the score.

A sister residue may be fixed in its own clade because it works against that clade’s background, in which case transplanting it is unsafe and a predictor’s benign call is a false negative. Separating that from genuine tolerance requires the coupled partner positions, which is the computational epistatic rescue problem restated [49]. That framing predicts a split and not a direction. Sister states whose partners match should be benign-enriched and those whose partners differ should not be. The directional variant tension framework supplies the per-direction baseline that a symmetric exchangeability matrix cannot represent, and is the natural next control [50]. Our estimate is also an association in curated data, not an experiment; the experiment is a deep mutational scan at clade-partitioned positions, scoring sister states against matched substitutions.

## Conclusions

A conservation score is a measurement of a position, a sequence set and a depth parameter, and the two failure modes characterised here are consequences of treating only the first as real. Gap inflation is definitional, arising from an alphabet that counts absence as a character, and an independent predictor identifies the gap-free recomputation as the better measure by a margin of 0.135 in rank correlation that replicates across two cohorts. Clade partitioning is structural, arising because paralogues diverge precisely at the positions that distinguish them, and it moves a position by about two grades on the nine-grade scale that users read while flipping 2.7 to 8.3 per cent of positions from variable to conserved, with a median across eight families of 6.6 per cent.

Gap inflation is removable by recomputation. Excluding the gap from the alphabet raises the AUC against the AlphaMissense-derived label from 0.832 to 0.919 at no cost in data or computation, and reporting the gap fraction alongside the score costs a column in a table. The tooling has existed in phylogenetics for twenty years in the form of column-trimming filters [14, 15, 17]; it simply has not been carried across to routine conservation reporting. Clade partitioning cannot be fixed by recomputation, because the information is genuinely absent from a pooled family alignment; it can only be flagged, and flagging it requires subfamily sampling that shallow alignments do not possess. A user who computes a conservation score from a default alignment can correct the first mode, and the panel states whether the second is testable at that depth, so two of its three columns are available wherever a score is computed and the third either carries a value or is reported as unavailable.

The limit is not a call for better software. The subfamily sampling needed to detect clade partitioning is roughly an order of magnitude deeper than the sampling a conservation score needs to be computed at all, and the alignments conservation scores are routinely computed from are rarely deep enough to expose the partition distorting them. Where the sampling is unavailable, the honest output is not a corrected score but an explicit statement that the score is unverified in that direction.

Computational evidence enters variant classification through the PP3 and BP4 criteria, and the recent recalibration of those criteria assigns evidence strengths on the basis of predictor scores whose own inputs include conservation [51, 52, 53]. A per-position reliability flag is exactly the kind of qualifier that framework can consume. A benign-side call resting on a column that is nine tenths gap, or a tolerance call resting on a column whose variation is entirely between paralogues, is not evidence of the strength the score implies.

The panel has been applied to the pI-code design gate, where it replaces an unqualified pass on four candidate substitutions with explicit verdicts, two marginally diagnostic and two indeterminate against the tree-aware null, and leaves a stricter gate than the three-paralogue test it replaces (Table 1). The gap correction was applied the same way to our own published conserved-residue scores, where it raises the rigid score’s discrimination in the human cohort from AUC 0.832 to 0.919 and places two of that paper’s four nominations on columns a 50 per cent gap filter excludes.

The sister-state prediction is tested here on 1,565 curated variants and remains open to a larger test at scale the moment a large enough set of clade-partitioned positions with curated variants is assembled, and the directional and epistatic controls it requires are already specified [49, 50]. More broadly, a conservation reliability panel is a natural covariate for variant-effect benchmarking. Existing benchmarks stratify by protein, by assay, by taxon and by alignment depth, but not by the per-position gap fraction or by clade partitioning [54, 55], and stratifying a benchmark that way would show whether predictors inherit these artefacts or merely tolerate them.

The broader argument is one about reporting and not about any single statistic. The conservation scores in common use are computed from alignments whose depth and composition are set by defaults and then largely discarded, and the score is delivered as though it were a property of the protein. Reporting the gap fraction, the effective comparator count and a tree-corrected partition statistic alongside the grade costs two numbers the computation already holds and one, the partition statistic, that costs a tree and a per-column simulation and is unavailable at shallow depth. Where a score is used to decide that a residue may be moved, or that a variant is tolerated, those three numbers say whether the score is entitled to the weight being placed on it.

## Methods

### Family assembly and the orthology criterion

The peroxidase family is the 80 reviewed members of Pfam PF03098 [56, 57] retrieved from UniProt [58], labelled into the four mammalian-type subfamilies by protein name, giving 6 lactoperoxidases including human, 5 thyroid peroxidases, 4 eosinophil peroxidases and 2 myeloperoxidases, with the remaining 63 family members unlabelled and excluded. The syndecan, nectin, junctional adhesion molecule, Eph, connexin, sodium channel alpha subunit and fibroblast growth factor receptor families were assembled from the EBI Proteins API. Orthologues are entries whose primary gene name equals the requested symbol, case-insensitively, within Mammalia as judged from the organism lineage. Entries matching a symbol only as a synonym are rejected by re-checking the primary name; second entries for the same gene and organism are dropped and listed; and a length filter dropping any sequence whose length differs by more than 40 per cent from that gene’s own human anchor removes fragments. Reviewed entries are fetched separately and always retained, because a single combined page placed the human F11R entry past the pagination limit. Subfamilies are capped at 20 or 30 sequences, reviewed first. Every excluded entry is written to a CSV with its reason. The clade restriction carries weight, since gene symbols collide across kingdoms: a search for SDC1 returns a Schizosaccharomyces pombe Set1-complex subunit. The criterion was chosen because every admission and rejection it makes is recorded and checkable, and because it can be held against the reciprocal-best-hit and tree-reconciliation criteria that dedicated resources apply [59, 60]; the orthologue-paralogue distinction is itself contested as a predictor of functional equivalence [61, 62], so the assembled families are validated against reconciled gene trees instead of the criterion being assumed sound.

### Alignment, position mapping and the subfamily-explained fraction

Sequences are aligned with FAMSA under a UPGMA guide tree [63]. Reference positions are mapped to alignment columns through the gapped reference row, which is required to reconstruct its input sequence exactly before any statistic is computed. For a column, with X the residue over the comparison set and Y the subfamily label, the subfamily-explained fraction is

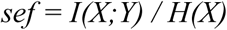

the mutual information normalised by the column entropy, which is the uncertainty coefficient of X given Y. This is the statistic of SDPpred [25] and is adopted unchanged. Normalising by H(X) instead of by min(H(X), H(Y)) or by the geometric mean keeps the denominator independent of the subfamily sizes. The reference sequence is excluded from the comparison set, so an identity is the fraction of the other sequences carrying the reference residue. Columns below an entropy floor of 0.5 bits are reported as conserved and carry no value, because the ratio is unstable as H(X) approaches zero; subfamilies with fewer than two ungapped sequences at a column are excluded from it; and columns above a gap fraction of 0.25 are not assessed, 0.25 being the level above which a column is not reliably alignable. The panel reports no estimate in either direction below five effective comparators at the column. The rebuild of the published pipeline in the previous section filters instead at a gap fraction of 0.50, the level at which a column holds more gap than residue, because that is the filter the comparison with the published score requires; the two thresholds govern different analyses and are not alternatives.

### Null models

The label-permutation null permutes subfamily labels across sequences, holding the residue composition and the subfamily sizes fixed, over 1,000 permutations throughout and 2,000 in the positive control, and the chance-corrected value is

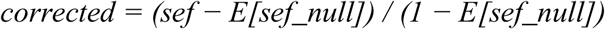

floored at zero. The tree-aware null builds a neighbour-joining tree [43] from Poisson-corrected p-distances computed over positions ungapped in both sequences, with negative branch lengths clamped to zero and counted. Characters are simulated down the tree under F81 [44] with equilibrium frequencies set to the column’s own composition, with transition probability

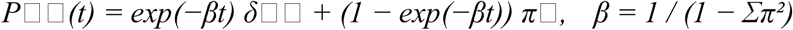

so that one unit of branch length is one expected substitution. Branch lengths are scaled per column by a multiplier found by bracketing on a 16-point log-rate grid and then bisecting eight times, chosen so that the mean Fitch parsimony score [45] of 80 simulated replicates at each grid point matches the observed one within a tolerance of the greater of 0.5 substitutions and 5 per cent of the score. The reported null is 1,000 replicates at the calibrated rate. A column is called diagnostic at a chance-corrected value of 0.50 or above with an empirical p of 0.05 or below, tolerant below 0.25, and indeterminate otherwise.

The tree is neighbour joining and not maximum likelihood, and the per-subfamily cap makes subfamilies balanced by construction. Both are declared parameters, and a better-specified tree would move only the tree-aware arm and towards a stricter null.

### Interval estimation and resampling

Every interval quoted in this work is a 95 per cent interval. The one on the Mantel-Haenszel odds ratio is the Robins-Breslow-Greenland interval, computed from the same family-by-conservatism strata the estimator uses. The intervals on areas under the curve are percentile bootstraps over the scored residues at a fixed seed, so a published bound reproduces exactly. A percentile bound is itself a Monte Carlo estimate, and the replicate count is set by how close the bound lies to a decision. The areas measured against the AlphaMissense label are bootstrapped at 2,000 replicates, and the intervals they return, 0.032 and 0.019 wide, lie far enough from each other and from any threshold that the count does not affect what they support. Everything measured against the clinical labels uses 50,000, because the paired difference there has a lower bound near zero. At 2,000 replicates that bound covers −0.002 to +0.008 across thirty seeds, so whether the interval excluded zero would depend on which seed was published; the run reports that sweep beside the interval it publishes. Two areas measured on one set of residues are paired, and their difference is bootstrapped by resampling residues once per replicate and recomputing both areas on that resample, which removes the shared sample from the comparison; the marginal intervals of such a pair overlap even where the difference is determined, so the difference is the quantity reported.

### Clinical labels

The clinical comparison uses ClinVar germline classifications [48] as a label independent of AlphaMissense, which is otherwise both the arbiter between the two conservation definitions and the source of the label in the published-score reanalysis. Pathogenic, likely pathogenic and the combined call are taken as positive, benign, likely benign and the combined call as negative, and every other value, uncertain significance and conflicting classifications among them, is dropped. A residue carrying calls on both sides would be dropped as conflicting; none does. That leaves 86 residues of the human cohort, 34 positive and 52 negative, and the four pipeline variants are scored against them unchanged. ClinVar is not redistributed; the derived labels and the resulting areas are deposited.

### Depth ladder and conservation grades

The four comparator sets are drawn from the single peroxidase source file so that sequences are identical between depths, and each depth is aligned independently with its own reference-row position map. Conservation grades are computed as equal-count percentile ranks of identity within each depth, in nine bins, which reproduces the within-protein normalisation ConSurf applies [2] and makes grades comparable across depths without reference to absolute divergence. An inversion is defined as a family-depth identity below 0.50 together with an own-subfamily identity of 0.90 or above.

### Gap-free recomputation and analysis of published scores

Gap-free conservation renormalises the twenty residue frequencies to the ungapped mass and returns

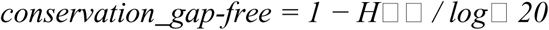

where H is the entropy over the twenty residue symbols alone. The published pipeline was rebuilt in four variants, with normalised dynamics recomputed per protein over whichever residues each variant retains and both scores min-max rescaled to 0 to 10 per protein as published [11]. Discrimination is measured against that paper’s own label, a mean AlphaMissense score above 0.90 as positive and below 0.20 as negative; this measures agreement with AlphaMissense and not independent accuracy, and no claim of outperforming AlphaMissense is made on it. The area under the curve is computed from ranks as the Mann-Whitney statistic. Correlations throughout are Pearson on ranks, and partial correlations are the rank-based partial form.

### Structural mapping

The lactoperoxidase fold in Figure 4 is PDB entry 7DE5, chain A, mapped to the human UniProt numbering by pairwise alignment of the deposited chain against the human sequence. The deposited chain is yak lactoperoxidase, so the map is cross-species and carries 595 residues at 85 per cent identity, which is sufficient for a positional correspondence and is not used for any structural measurement. Every catalytic and cofactor position of the ten-position control in Table 1 was confirmed to carry its expected residue in the mapped structure before plotting. Backbone alpha carbons are projected onto their two leading principal axes and rendered in depth order.

### Halting conditions and verification

Every analysis script halts rather than warning and writes output only after all checks pass, with Table S5 listing the condition each one halts on. A separate control script recomputes every quoted control number from the source FASTA without importing the analysis implementation, and runs 26 checks, all passing.

## Data and software versions

Python 3 with numpy and pandas, pyfamsa for FAMSA alignment and matplotlib for figures. UniProt sequences were retrieved from the EBI Proteins API. Curated human variants are the UniProt humsavar index, release 2026_03, 85,015 single amino acid variants [58]. AlphaMissense scores are those assembled in our previous work [10, 11]. All analysis scripts, the assembled families with their exclusion lists, and the per-position outputs with their run logs are provided.

## Supplementary information

Supplementary Information contains Supplementary Notes S1 to S3, Supplementary Tables S1 to S8 and Supplementary Figures S1 to S4. Note S1 gives the entropy-matching calibration argument in full, including why the target is unreachable and what the symptom looks like; Note S2 the orthology criterion and what it can establish; and Note S3 what stratification does to the sister-state estimate. Supplementary Tables S10 to S19 are the long per-position tables, 52,454 rows in all, in the accompanying workbook Supplementary_Tables.xlsx: the tree null and the reliability panel for every family, the curated variants with their sister-state annotation, the depth ladder, the positive-control sweep, per-protein gap summaries, the sister-state strata and every sequence rejected during family assembly with its reason.

## Code and data availability

The thirteen analysis scripts written for this work, the per-position output tables, the assembled family alignments with their exclusion lists and the raw run log of every result quoted here are at https://github.com/karagol-taner/conservation-score-reliability. Every figure rebuilds from the deposited tables alone, and the deposit includes the control verifier, which recomputes the positive control from the source alignment without importing the implementation of the statistic it checks, sharing only the FASTA reader, the aligner and the subfamily labels. A browser tool at https://karagolresearch.com/gapcheck, mirrored at https://tanerkaragol.com/gapcheck, applies the panel’s gap checks to an uploaded alignment and reports for each column the gap fraction, the effective comparator count and the gap-free recomputation. It counts each sequence once over the columns as supplied, so its values need not match a frequency file that weights sequences or indexes positions against a target sequence, and no number reported in this work comes from it. The EVcouplings frequency files and AlphaMissense scores are those assembled in our previous work and are cited; source sequences are public in UniProt and the humsavar release is distributed by UniProt. The ClinVar classifications used for the clinical comparison are cited [48] and not redistributed; the 86 residue-level labels derived from them, and the areas under the curve computed from those labels, are in the deposit so that comparison reproduces from it alone. No new sequence or structure data were generated.

## Competing financial interests

Both authors are inventors on a patent application for the pI-code design method, which gates candidate substitutions on a conservation test of the kind this paper evaluates [34]. Both authors therefore have an interest in that method, and the following safeguards apply. No designed protein is produced or experimentally evaluated here and no claim is made about the method’s performance; the four candidate positions appear as a worked example and are re-gated against the panel in Table 1. Two directional predictions were registered in the analysis scripts before the numbers existed and both are reported as they came out: that alignment depth would predict gappiness, which it did not at a rank correlation of −0.034, and that variants to a sister subfamily’s fixed state would be enriched for benign classification, which held across all sister-state variants at a one-sided p of 0.0012 and reached a one-sided p of 0.070 in the narrower registered arm, short of the 0.05 threshold. Separately, A.K. and T.K. hold further invention disclosures and pending patents spanning two technology areas unrelated to this study: macromolecular structures for neurological indications; responsive biomaterials for microenvironment-targeted delivery. A.K. is an incoming Novo Nordisk AI Fellow at MIT, starting November 2026. T.K. holds an Anthropic AI for Science award for work unrelated to this study, whose use here is declared under Funding. Beyond that, this research received no financial or operational support from these entities, and none from the intellectual property or fellowship funding named above.

## Funding

No grant funding was received for this work. The large language model assistance declared below used model credits from an Anthropic AI for Science award held by T.K. for unrelated work. Anthropic had no role in the design of the study, the analysis, the interpretation of the results or the decision to publish.

## Ethics approval

Not applicable. This work analyses public datasets and generated no new data from human participants or animals.

## Consent to participate

Not applicable.

## Consent for publication

Not applicable.

## Author contributions

T.K. conceived the study, wrote the analysis code, performed the analysis and wrote the manuscript. A.K. contributed to the design of the analysis and to the interpretation of the results, and revised the manuscript. Both authors read and approved the final manuscript.

## Declaration of generative AI in the writing process

Large language model (Claude by Anthropic, Opus) assistance was used for code development and manuscript preparation (language editing). All analyses, numbers and conclusions were produced by the scripts described in the Methods and verified by the authors against their recorded output.

## Notes

https://github.com/karagol-taner/conservation-score-reliability

