## Supplementary Information for "Two opposite failure modes in alignment-based conservation scores, one inflating and one concealing constraint, are computable before the score is used"

This file contains Supplementary Notes S1 to S3, Supplementary Tables S1 to S9 and Supplementary Figures S1 to S4. The long per-position tables, Supplementary Tables S10 to S19, are in the accompanying workbook Supplementary_Tables.xlsx, whose first sheet lists their contents and the script that produced each.

### Table of Contents


| Item | Title | Page |
|---|---|---:|
| Note S1 | Why the tree-aware null is calibrated on parsimony and not on entropy | 3 |
| Note S2 | The orthology criterion and what it can establish | 4 |
| Note S3 | What stratification does to the sister-state estimate | 5 |
| Table S1 | Family assembly: sequences kept and rejected | 6 |
| Table S2 | Conservation by decile of column gap fraction | 7 |
| Table S3 | The ten commonest ordered sister substitutions | 8 |
| Table S4 | Family-by-conservatism strata of the sister-state test | 9 |
| Table S5 | Halting conditions by script | 10 |
| Table S6 | Clade-partitioning rate across nine family runs | 11 |
| Table S7 | Reanalysis of the published conserved scores | 12 |
| Table S8 | Sister-state substitutions against all others, by family | 13 |
| Table S9 | Discrimination against ClinVar germline classifications | 14 |
| Figure S1 | The calibration diagnostic for the tree-aware null | 15 |
| Figure S2 | Where each null places its expectation | 16 |
| Figure S3 | Alignment depth does not predict gappiness, which is why the two failure modes need separate flags | 17 |
| Figure S4 | What the gap artefact did to a published derived score | 18 |

**Note S1. Why the tree-aware null is calibrated on parsimony and not on entropy**

The tree-aware null simulates a neutral character along the family's own tree and asks whether the observed column is more clade-partitioned than that. The simulation needs a rate, and the rate has to be calibrated so that the simulated character is as variable as the observed one. The obvious quantity to match is entropy, searching for the branch-length multiplier at which the mean simulated column entropy equals the observed column entropy. Entropy cannot serve, because of how the plug-in estimator behaves on a finite number of tips.

The equilibrium frequencies of the F81 process are set to the column's own composition, which is deliberate and is what isolates the arrangement of residues among clades from which residues they are. A consequence is that at high rate the tips become independent draws from that composition. The mean plug-in entropy of n such draws sits below the entropy of the composition itself by the finite-sample bias, approximately $(k - 1)/(2n \ln 2)$ bits for k occupied states, which is about 0.27 bits for a seven-state column over sixteen sequences. The observed column's plug-in entropy is computed from that same composition without the correction. The target therefore sits above anything the simulation can reach, the search runs to the top of the rate grid, and at the top of the grid no phylogenetic correlation survives. The tree null silently becomes the permutation null.

A null whose target is unreachable reads as a careful control confirming that ancestry does not matter much. Every column whose target cannot be reached is reported as a calibration failure, and that rate is the only symptom, which is easy to dismiss as numerical awkwardness. Figure S1a shows the rate under parsimony matching, 0.6 per cent on the peroxidases and 17.8 per cent on the Ephs.

Fitch parsimony is the quantity the calibration matches. It rises monotonically from zero as the rate rises, it is reachable by construction, and it is a count of changes on the tree and not a property of the tip distribution, so the finite-sample argument does not apply to it. Figure S1b shows the match achieved: the worst deviation between observed and mean simulated parsimony over the calibrated peroxidase columns is 0.46 substitutions. On that family the clade-partitioning rate is 7.5 per cent against the permutation null's 68.3.

**Note S2. The orthology criterion and what it can establish**

The orthology criterion decides which sequences enter a family, so the clade-partitioning rate is a statement about the set the criterion admits.

The criterion is a UniProt entry whose primary gene name equals the requested symbol, case-insensitively, within Mammalia as judged from the organism lineage, one sequence per gene and organism, with a per-gene length filter at 40 per cent of that gene's own human anchor. It is a weaker relation than the reciprocal-best-hit and tree-reconciliation criteria dedicated resources apply, which is why the assembled families are audited against Ensembl Compara below.

A gene symbol is not unique across clades. A search for SDC1 returns a Schizosaccharomyces pombe Set1-complex subunit alongside the syndecans, which the clade filter excludes.

Gene lengths differ within a family, and syndecan-3 is 442 residues against syndecan-1's 310, so the length filter is anchored on each gene's own human sequence and not on the family reference, where a 40 per cent band drawn around the latter would exclude every curated SDC3 entry.

A single combined request can place a reference sequence past the page limit, as it does for human F11R in the JAM family, so reviewed entries are fetched separately and always retained.

What the criterion cannot establish is one-to-one orthology. A lineage-specific duplication that retains the ancestral gene symbol will pass. The deep families rest on 10 to 18 curated entries out of 90 to 280, so a proportion of the remainder will be misassigned. The sevenfold sensitivity of the clade-partitioning rate to sampling depth, reported in the main text, is the quantitative form of that concern, and every rejected sequence is listed with its reason in sheet S18 of the workbook so the criterion can be audited rather than taken on trust.

Ensembl Compara, whose orthology calls come from reconciled gene trees, adjudicates 765 of the 1,102 non-human sequences in the eight families. Of those 765, 757 are one-to-one orthologues of the human gene they were matched to, 99.0 per cent, with per-family rates from 98.3 per cent in the Eph receptors to 100 per cent in the peroxidases and connexins. The eight exceptions are orthologues that are not one-to-one, in goat, wombat, rat, kangaroo rat, chimpanzee and brown bear, with goat and kangaroo rat contributing two each, which is the lineage-specific duplication described above. Of the 337 sequences Compara does not adjudicate, 307 are in species for which it holds no genome and 30 are non-mammalian members of the Pfam-derived peroxidase file with no human counterpart in it. The check is independent of every result reported here, and over the 765 sequences it adjudicates, 1.0 per cent fall outside a one-to-one relation.

**Note S3. What stratification does to the sister-state estimate**

The sister-state test asks whether human missense variants to a residue that a sister subfamily carries in the equivalent position are enriched for benign classification. Two confounds make a pooled comparison meaningless, and both are handled by stratification and not by argument.

The first is that sister states are disproportionately conservative exchanges. Across the eight families 3,063 positions carry at least one sister state, giving 8,649 ordered substitution instances in 361 distinct classes; the ten commonest are listed in Table S3 and every one of them is a conservative exchange. A test that ignored this would partly rediscover that conservative substitutions are benign.

The second is ascertainment. A disease gene is sequenced in patients, so its curated variants are overwhelmingly pathogenic. The benign share among variants at these positions runs from 0.084 in the connexins to 0.762 in the Eph receptors, a ninefold difference that has nothing to do with the substitutions being compared. Pooling the families into one table compares families.

Mantel-Haenszel conditions on each stratum's own margins, so a family is compared only against itself and a conservative exchange only against other conservative exchanges. Table S4 gives the sixteen family-by-conservatism strata. Nine carry information; the other seven have an empty margin, contribute nothing to either side of the estimator, and are shown with their counts so the reader can see which they are.

The unstratified odds ratio is 4.42 and the stratified one is 2.46, so more than half the naive effect was the difference between families. Restricting further to positions the tree-aware null calls diagnostic, the restricted form in which the prediction was registered, gives 2.07 at a one-sided p of 0.070. We treat the unrestricted arm as primary because the mechanism is the sister state itself, a residue the fold is observed to accept in a related clade, and that property holds at a position whatever a null later says about it. Both arms are reported with their counts, so the restriction can be applied or not by the reader.

### Supplementary Tables

**Table S1. Family assembly. Sequences kept after the clade filter, the primary-gene-name recheck, the one-per-gene-and-organism rule and the per-gene length anchor, against the number rejected. Every rejection is listed with its reason in sheet S18 of the workbook. The peroxidase family is not listed here because it is the 80 reviewed members of Pfam PF03098 rather than an assembled set.**

| Family | Assembled file | Sequences kept | Rejected |
|---|---:|---:|---:|
| syndecans (reviewed only) | syndecans_reviewed.fasta | 18 | 18 |
| syndecans | syndecans_all.fasta | 120 | 2,289 |
| nectins | nectins_all.fasta | 120 | 2,287 |
| JAMs | jams_all.fasta | 90 | 1,971 |
| FGFRs | fgfrs_all_all.fasta | 120 | 3,892 |
| connexins | gjbs_all_all.fasta | 150 | 2,185 |
| sodium channel alpha subunits | scns_all_all.fasta | 200 | 7,071 |
| Eph receptors | ephs_all.fasta | 280 | 10,934 |

**Table S2. Mean conservation by decile of column gap fraction, over the 55,816 columns of the dynamics-aware cohort, as reported and recomputed over the twenty residue symbols renormalised to the ungapped mass.**

| Gap fraction of the column | Columns | Conservation as reported | Conservation gap-free |
|---|---:|---:|---:|
| 0 to 10% | 27,890 | 0.405 | 0.418 |
| 10 to 20% | 7,464 | 0.369 | 0.409 |
| 20 to 30% | 3,920 | 0.349 | 0.368 |
| 30 to 40% | 3,097 | 0.365 | 0.341 |
| 40 to 50% | 2,087 | 0.416 | 0.342 |
| 50 to 60% | 1,632 | 0.474 | 0.326 |
| 60 to 70% | 1,994 | 0.580 | 0.394 |
| 70 to 80% | 2,054 | 0.655 | 0.361 |
| 80 to 90% | 2,189 | 0.776 | 0.415 |
| 90 to 100% | 3,489 | 0.934 | 0.597 |

**Table S3. The ten commonest ordered sister substitutions across the eight families, out of 8,649 instances in 361 distinct classes at 3,063 positions. The ten together account for 16.8 per cent of instances and every one is a conservative exchange, which is why the sister-state test is stratified on substitution conservatism.**

| Ordered substitution | Instances | Share of all instances |
|---|---:|---:|
| V → I | 186 | 2.2% |
| I → V | 180 | 2.1% |
| T → S | 159 | 1.8% |
| A → S | 143 | 1.7% |
| I → L | 140 | 1.6% |
| D → E | 139 | 1.6% |
| E → D | 139 | 1.6% |
| K → R | 136 | 1.6% |
| S → A | 119 | 1.4% |
| V → L | 115 | 1.3% |

**Table S4. The 16 family-by-conservatism strata of the sister-state test, unrestricted arm. 9 carry information; the rest have an empty margin and contribute nothing to the Mantel-Haenszel estimator.**

| Family | Stratum | Sister benign / total | Other benign / total | Informative |
|---|---:|---:|---:|---:|
| Peroxidases | conservative | 1 / 1 | 11 / 16 | yes |
| Peroxidases | non-conservative | 1 / 1 | 18 / 35 | yes |
| Syndecans | conservative | 0 / 0 | 4 / 4 | empty margin |
| Syndecans | non-conservative | 1 / 1 | 2 / 2 | empty margin |
| Nectins | conservative | 0 / 0 | 1 / 1 | empty margin |
| Nectins | non-conservative | 0 / 0 | 1 / 2 | empty margin |
| Junctional adhesion molecules | conservative | 0 / 0 | 0 / 1 | empty margin |
| Junctional adhesion molecules | non-conservative | 0 / 0 | 1 / 4 | empty margin |
| Fibroblast growth factor receptors | conservative | 1 / 1 | 9 / 29 | yes |
| Fibroblast growth factor receptors | non-conservative | 0 / 0 | 19 / 181 | empty margin |
| Connexins | conservative | 1 / 3 | 6 / 58 | yes |
| Connexins | non-conservative | 2 / 3 | 16 / 204 | yes |
| Sodium channel alpha subunits | conservative | 9 / 31 | 42 / 208 | yes |
| Sodium channel alpha subunits | non-conservative | 4 / 23 | 57 / 653 | yes |
| Eph receptors | conservative | 14 / 15 | 20 / 24 | yes |
| Eph receptors | non-conservative | 7 / 8 | 41 / 56 | yes |

**Table S5. Conditions on which each script halts without writing output. Every analysis script halts rather than warning, because a warning in a log is not read in time to stop a run. The conservation-recovery condition, which requires the reported conservation column to be reproduced by $1 - H/\log_2 21$ before anything downstream runs, is what establishes that the gap artefact is definitional and not statistical.**

| Script | Condition that halts the run |
|---|---|
| partition.py | A gapped reference row that does not reconstruct its input sequence; a sequence matching two subfamily patterns, so the partition is not a partition; an orthologue set overlapping the paralogue set |
| 01, 02, 05, 07 | An alignment or comparator depth below its stated sequence count; a subfamily left with no members |
| 01 | A catalytic or cofactor control residue failing in either direction |
| 03, 04, 06 | A frequency file lacking residue columns; a gap column outside the unit interval; a conservation column not reproduced by $1 - H/\log_2 21$ |
| 05, 06 | A curated variant whose wild-type residue disagrees with the sequence it is mapped onto |
| 07 | A calibration target unreachable across the whole rate grid, counted and reported per column |
| 08 | An entry matching the requested symbol only as a synonym; an entry outside the requested clade; a second entry for the same gene and organism |
| 09 | A variant table missing a required column; a classification outside the humsavar vocabulary; zero sister-state variants pooled, which would mean the join failed |
| build_figures.py | A missing tree cache; a tree cache holding the wrong number of tips |
| make_supplementary_tables.py | A missing source file; an empty sheet; a column set differing between two files being concatenated |
| verify_manuscript.py | Any quoted number not matching its source; a citation out of first-mention order; an uncited reference |

**Table S6. Clade-partitioning rate across nine family runs under both null models. Columns tested is the number the tree null returned a non-conserved call for, which is the denominator of both percentages. Figure 5b plots the two rates.**

| Family | Sequences | Per subfamily | Columns tested | Permutation null | Tree-aware null |
|---|---:|---:|---:|---:|---:|
| Peroxidases (PF03098) | 17 | 6/5/4/2 | 492 | 68.3% | 7.5% |
| Syndecans, reviewed only | 18 | 6/4/3/5 | 146 | 85.6% | 4.1% |
| Syndecans | 120 | 30 each | 161 | 93.8% | 29.8% |
| Nectins | 120 | 30 each | 429 | 90.7% | 30.5% |
| Junctional adhesion molecules | 90 | 30 each | 227 | 84.1% | 38.3% |
| Connexins (GJB) | 150 | 30 each | 137 | 96.4% | 32.8% |
| Sodium channel alpha subunits | 200 | 20 each | 1,160 | 97.6% | 49.7% |
| Fibroblast growth factor receptors | 120 | 30 each | 421 | 88.8% | 58.4% |
| Ephs | 280 | 20 each | 725 | 100.0% | 69.1% |

**Table S7. Reanalysis of the published rigid and dynamic conserved scores, four pipeline variants. Variant A reproduces the published pipeline, at a correlation between the two scores of 0.080 against the published 0.08 and with raw conservation at 0.903 against 0.91. Figure S4a plots the areas under the curve.**

| Variant | ρ(DCS, RCS) | ρ(RCS, conservation) | AUC RCS | AUC DCS | Top-decile DCS n | Gappy share |
|---|---:|---:|---:|---:|---:|---:|
| A, as published | 0.080 | 0.903 | 0.832 | 0.438 | 1,024 | 44.7% |
| B, gap filter | −0.093 | 0.888 | 0.899 | 0.492 | 863 | 0.0% |
| C, gap-free conservation | 0.060 | 0.919 | 0.919 | 0.531 | 1,024 | 28.0% |
| D, both | −0.044 | 0.902 | 0.916 | 0.541 | 863 | 0.0% |

**Table S8. Sister-state substitutions against all others, by family. Counts are curated human missense variants mapped to the family reference and classified as benign-side (LB/B) or pathogenic-side (LP/P); unclassified variants are excluded. The pooled row is the sum and not the estimate: the stratified Mantel-Haenszel odds ratio is 2.46 with a one-sided p of 0.0012. Figure 6a plots the percentages.**

| Family | Sister-state benign/total | Other benign/total | Sister benign % | Other benign % |
|---|---:|---:|---:|---:|
| Peroxidases | 2/2 | 29/51 | 100.0% | 56.9% |
| Syndecans | 1/1 | 6/6 | 100.0% | 100.0% |
| Nectins | 0/0 | 2/3 | - | 66.7% |
| Junctional adhesion molecules | 0/0 | 1/5 | - | 20.0% |
| Eph receptors | 21/23 | 61/80 | 91.3% | 76.2% |
| Fibroblast growth factor receptors | 1/1 | 28/210 | 100.0% | 13.3% |
| Connexins | 3/6 | 22/262 | 50.0% | 8.4% |
| Sodium channel alpha subunits | 13/54 | 99/861 | 24.1% | 11.5% |
| Pooled | 41/87 | 248/1,478 | 47.1% | 16.8% |

**Table S9. Discrimination against ClinVar germline classifications, over the 86 residues of the dynamics-aware human cohort that carry an unambiguous classification, 34 pathogenic and 52 benign. All four pipeline variants and both published scores are listed. The main text quotes the two rigid-score rows for variants A and C. Intervals are percentile bootstraps over the scored residues at 50,000 replicates and a fixed seed. The gap-free dynamic score is the one of the eight areas whose interval clears 0.5, which holds against this label and not against the AlphaMissense label of Table S7, where the same score stays at chance.**

| Variant | Score | Clinical AUC[1] | 95% interval | Residues |
|---|---|---:|---:|---:|
| A, as published | RCS | 0.535 | 0.403 to 0.665 | 86 |
| A, as published | DCS | 0.555 | 0.424 to 0.687 | 86 |
| B, gap filter | RCS | 0.588 | 0.448 to 0.725 | 71 |
| B, gap filter | DCS | 0.494 | 0.354 to 0.634 | 71 |
| C, gap-free conservation | RCS | 0.610 | 0.482 to 0.732 | 86 |
| C, gap-free conservation | DCS | 0.639 | 0.514 to 0.759 | 86 |
| D, both | RCS | 0.597 | 0.458 to 0.731 | 71 |
| D, both | DCS | 0.531 | 0.390 to 0.673 | 71 |
| C minus A, paired difference | RCS | +0.075 | +0.003 to +0.149 | 86 |

[1]The final row is the difference between the two rigid-score rows for variants C and A, which are measured on one set of residues. Each replicate resamples those residues once and recomputes both areas on that resample, so the shared sample cancels out of the difference. The interval on that row is on the difference and not on either area.

### Supplementary Figures

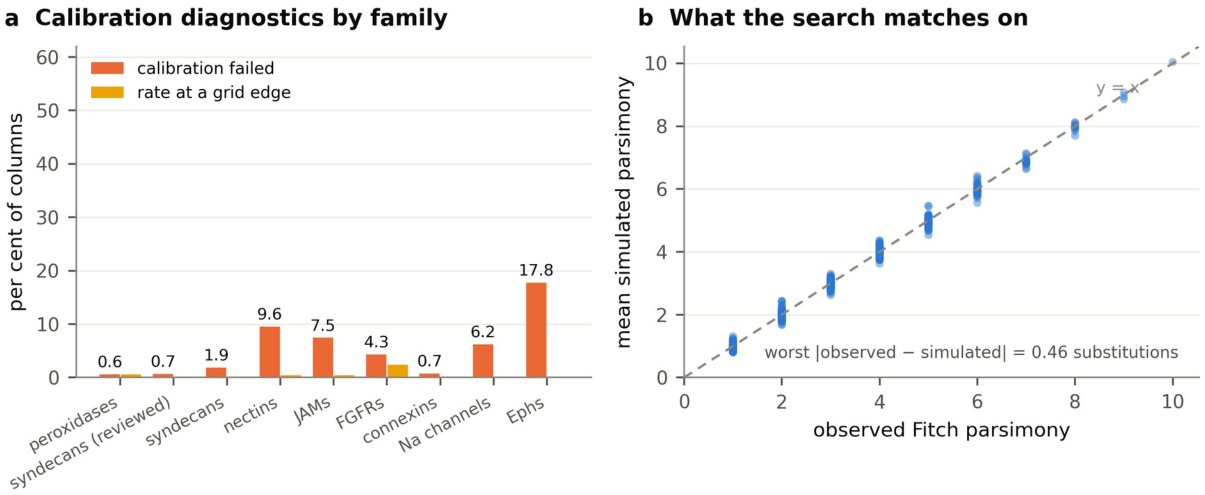


**Figure S1. The calibration diagnostic for the tree-aware null.**
(a) Percentage of columns whose per-column rate search failed to meet its parsimony target, and percentage whose fitted rate sat at an edge of the rate grid, by family. A null calibrated on a target it cannot attain fails on every such column, so this rate is the symptom to watch, and Note S1 gives the argument. (b) Observed Fitch parsimony against the mean parsimony of the simulated replicates, over the calibrated peroxidase columns. The search matches on this quantity, and the worst deviation is 0.46 substitutions.

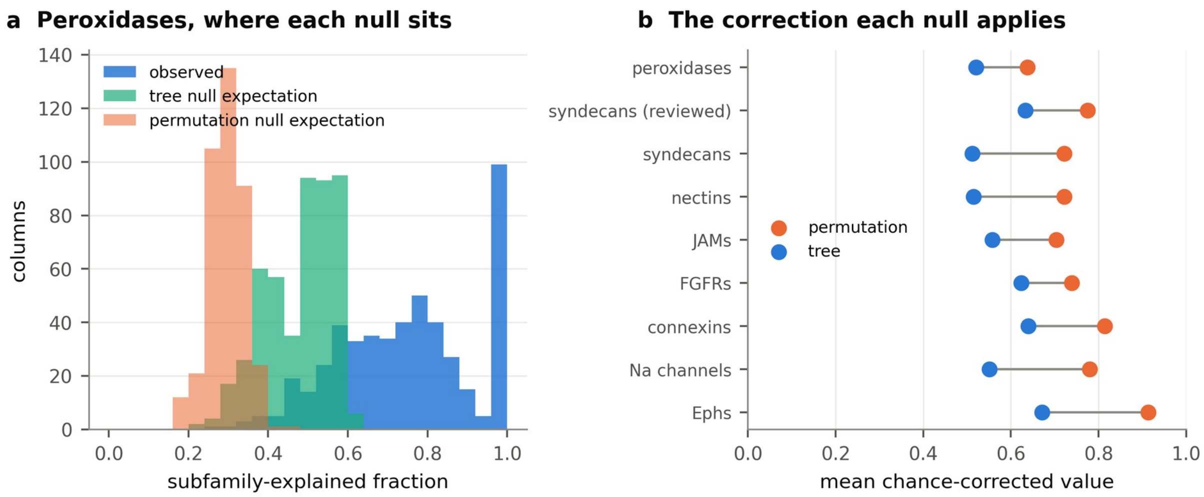


**Figure S2. Where each null places its expectation.**

(a) Distribution of the subfamily-explained fraction over the calibrated peroxidase columns: the observed values, the mean under the tree-aware null, and the mean under the label-permutation null, recovered from the chance-corrected values. The permutation null's expectation sits far below the observed values at almost every column, which is why it calls two thirds of them diagnostic. (b) Mean chance-corrected value under each null, by family. The gap between the two points is the correction the tree null applies and the permutation null does not.

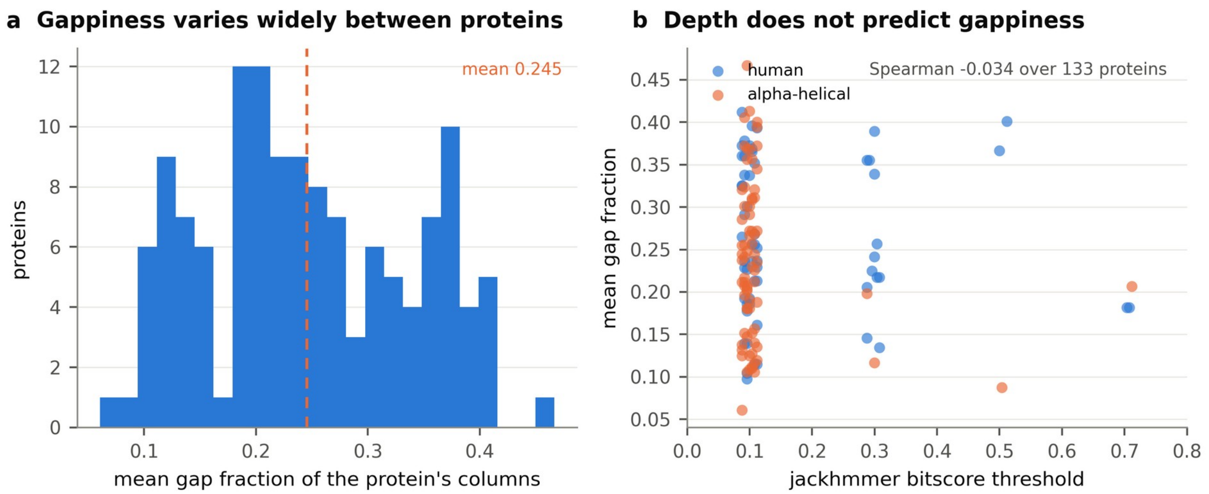


**Figure S3. Alignment depth does not predict gappiness, which is why the two failure modes need separate flags.**

(a) Distribution of mean gap fraction across the 133 proteins of the dynamics-aware cohort. (b) The same quantity against the per-protein jackhmmer bitscore threshold that sets alignment depth. A permissive search admitting distant sequences that cover less of the query was registered as the predicted cause of gappiness. The rank correlation is −0.034, so the gap fraction cannot be inferred from the search threshold the alignment was built at, and has to be reported per column.

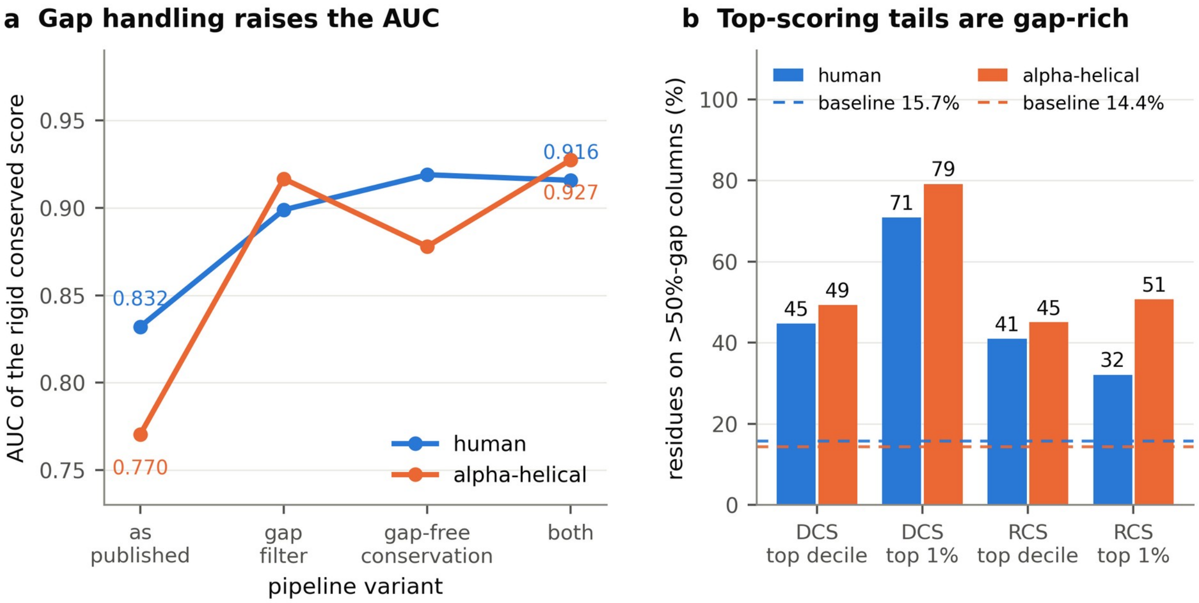


**Figure S4. What the gap artefact did to a published derived score.**
(a) Discrimination of our own published rigid conserved score against the published label, under four pipeline variants: as published, with gappy columns filtered, with gap-free conservation, and with both. Handling the gap raises the area under the curve from 0.832 to 0.919 in the human cohort and from 0.770 to 0.928 in the alpha-helical one, so the gap-inclusive alphabet limits the discrimination available to that pipeline. (b) Percentage of residues in the top decile and top percentile of each score that sit on columns more than half gap, over that paper's published score table and shown against the cohort base rate. The population described there as missed by standard conservation analysis is roughly half gap artefact at the decile and three quarters at the percentile, in both cohorts independently.
